# Unraveling the prevalence and multifaceted roles of accessory peptide deformylases in bacterial adaptation and resistance

**DOI:** 10.1101/2025.06.06.657839

**Authors:** Morgan Lambérioux, Magaly Ducos-Galand, Pierre-Alexandre Kaminski, Eloi Littner, Jean-Michel Betton, Ariel Mechaly, Ahmed Haouz, Didier Mazel

## Abstract

Peptide deformylases (PDFs) are enzymes that are essential for bacterial viability and attractive targets for antibiotic development. Yet, despite their conserved function, many bacteria encode multiple PDFs, a genomic feature whose prevalence and implications remain largely unexplored. Here, we reveal that nearly half of all bacterial genomes carry more than one PDF gene, frequently embedded within mobile genetic elements such as plasmids and integrons. In *Vibrio cholerae*, the accessory PDF (Def2_VCH_) confers reduced susceptibility to actinonin (ACT), the most studied PDF inhibitor, while still supporting bacterial growth in absence of the canonical PDF copies (Def1_VCH_). Crystallographic analysis shows that this reduced susceptibility stems from an arginine-to-tyrosine substitution that probably reduces ACT binding. Strikingly, this resistance signature is shared by integron-encoded PDFs, and transfer of an integron-encoded PDF cassette from *Pseudoxanthomonas* into a susceptible *V. cholerae* is sufficient to abolish ACT susceptibility. These findings expose a cryptic reservoir of resistance within the bacterial mobilome and highlight a challenge to the therapeutic potential of PDF-targeting antibiotics: resistance may not only emerge, but is already encoded, mobile, and ready to spread.

## Introduction

In bacteria, as well as in mitochondria and chloroplasts, translation is always initiated by a formylated methionine, which is usually cleaved during the translation (reviewed in (Giglione *et al*., 2015)). Peptide deformylases (PDFs) are metalloproteases that catalyze the removal of the formyl group from the initiator methionine, a necessary step in the N-terminal methionine excision (NME) process (Mazel *et al*., 1994; Giglione *et al*., 2015). PDFs are promiscuous enzymes with limited specificity, deformylating approximately 95% of the bacterial proteome (Bienvenut *et al*., 2015). Generally, only membrane or exported proteins, which have a hydrophobic signal peptide at their N-terminus, escape deformylation (Yang *et al*., 2022).

For over two decades, it has been observed that some bacteria possess multiple PDF encoding genes while they contain only one *fmt* gene, allowing the formylation of the initiator methionine (Margolis *et al*., 2000; Guilloteau *et al*., 2002). Notable species among these include important model organisms or pathogens such as *Staphylococcus aureus*, *Enterococcus faecalis*, *Bacillus subtilis*, *Streptococcus pneumoniae*, *Vibrio cholerae*, and *Pseudomonas aeruginosa* (Margolis et al., 2000). In certain cases, such as *S. aureus* and *S. pneumoniae*, substitutions in the conserved motifs essential for catalytic activity result in the inactivation of one of the PDFs in the laboratory conditions tested (Margolis et al., 2000; Margolis et al., 2001). In other examples, such as in *B. subtilis*, both PDFs are active and share similar enzymatic and inhibition properties (Haas *et al*., 2001). Despite the longstanding observation of multiple PDF genes, the evolutionary advantage of expressing several different PDFs remains unclear. Moreover, no large-scale genomic analysis has been performed to study the distribution of PDFs in bacterial genomes, to better understand their possible selective role.

Given their conservation and indispensable roles in the bacterial translation process, PDFs have been proposed as attractive targets for the development of new antibiotics (Mazel *et al*., 1994). Actinonin (ACT), a natural product made by bacteria from the genus *Streptomyces* (Gordon *et al*., 1962; Gordon *et al*., 1975), was identified as the first PDF inhibitor (Chen *et al*., 2000). Since then, numerous other PDF inhibitors have been discovered (reviewed extensively in (Jain *et al*., 2005; Sangshetti *et al*., 2015)). All peptide-based PDF inhibitors share a common mechanism of action: they bind to the catalytic pocket of the enzyme, thereby preventing substrate access and inhibiting its deformylase activity (Sangshetti *et al*., 2015). To date, three mechanisms of reduced susceptibility to ACT have been described: (1) drug export by eflux pumps (Chen *et al*., 2000; Margolis *et al*., 2000; Clements *et al*., 2001); (2) impairment of the formylation mechanism by inactivation of *fmt*, which directly catalyzes the formylation of initiator methionine, or of *folD* or *glyA*, which contribute to the synthesis of the formyl group donor essential for Fmt activity (Margolis *et al*., 2000; Nilsson *et al*., 2006; Duroc *et al*., 2009; Yang and Sun, 2016); and (3) overexpression or mutations of the PDF (Margolis *et al*., 2001; Dean *et al*., 2007). Although inactivation of the formylation process is the most common mechanism of reduced susceptibility, it significantly slows bacterial growth (Guillon *et al*., 1992; Mazel *et al*., 1994) and could lead to a drastic reduction in virulence, as shown for *S. aureus* in mouse models (Margolis *et al*., 2000), indicating that despite the development of resistance, PDF inhibitors may remain effective in treating bacterial infections due to the co-associated growth defect and reduced virulence.

PDFs are relatively small enzymes, typically ranging from 150 to 200 amino acids (aa) in prokaryotes and from 220 to 250 aa in eukaryotes. Despite sharing only 20–30% sequence homology between phylogenetically distant PDFs, all PDFs exhibit significant structural conservation (Giglione *et al*., 2004). They mainly differ in the structure of their C-terminal region, which, has been shown in *Escherichia coli* to facilitates interaction with the ribosome (Bingel-Erlenmeyer *et al*., 2008; Sandikci *et al*., 2013; Bhakta *et al*., 2019; Akbar *et al*., 2021). Most PDFs share a common catalytic core defined by three conserved sequence motifs (I, II, and III), which coordinate a metal ion and position key residues for substrate processing (Giglione *et al*., 2004).

Phylogenetically, PDFs are classified into four main group called types (Grzela et al., 2017). Type 1A and 1B PDFs, found in both eukaryotes and prokaryotes, have a characteristic C-terminal tail folded into an α-helix (Chan *et al*., 1997; Giglione *et al*., 2009). Type 2 PDFs are mostly found in Gram-positive bacteria and have a C-terminal tail folded into a β-strand (Giglione *et al*., 2009; Grzela *et al*., 2017). The structures of type 3 PDFs, as well as, of type 4 PDFs, have not yet been resolved. Type 3 PDFs, present in Archaea, protists, and some bacterial genera, display substitutions in conserved motifs I and II, which have been shown to lead to reduced enzymatic activity or even inactive enzymes (Margolis *et al*., 2000; Giglione *et al*., 2004; Bouzaidi-Tiali *et al*., 2007; Grzela *et al*., 2017). The type 4 PDFs remain less well-characterized, and are found in a broad variety of host including marine viral bacteriophage and eukaryotic parasite (Grzela *et al*., 2017).

The biological significance of encoding multiple PDF genes within the same genome remains poorly understood. One hypothesis is that this redundancy could be linked to resistance against PDF inhibitors, natural products synthesized by bacteria that may be encountered in diverse environmental contexts. With the rise of antibiotic resistance worldwide, it becomes urgent to identify new therapeutic targets. As PDF has been shown to be a promising target, understanding the distribution and role of these accessory PDFs is crucial. To address this knowledge gap, we explored the distribution of PDF across RefSeq bacterial genome and focused on the functional roles of some specific PDFs, including those found in *Vibrio* species. Our findings reveal that approximately half of the bacterial reference genomes possess multiple PDF genes, with significant intra-species variability. We discovered that PDFs can be encoded on mobile genetic elements (MGEs), such as plasmids and integrons, which likely contributes to this variability and suggests an adaptive role. Furthermore, our investigation of *Vibrio cholerae*, which encodes two type 1b PDFs with 50% sequence homology (Def1_VCH_ and Def2_VCH_), demonstrated that both PDFs are catalytically active. We found that Def2_VCH_ specifically confers reduced susceptibility to the natural PDF inhibitor, ACT. Additionally, the integron PDF cassette identified in a *Pseudoxanthomonas suwonensis* strain was shown to impart significant resistance to PDF inhibitors in both *V. cholerae* and *E. coli*. This resistance phenotype suggests that the dissemination of this cassette could pose a challenge for the therapeutic use of PDF inhibitors and raises questions about their viability in clinical applications.

These discoveries underscore the diverse distribution of PDF genes in bacteria and support the hypothesis that many additional PDF gene copies have been selected for their role in resistance to natural PDF-targeting antibiotics, thereby questioning the potential for clinical application of PDF inhibitors in treating bacterial infections.

## Materials & methods

### Bacterial Strains, Plasmids, and Primers

All bacterial strains and plasmids in this study are summarized in Supplementary **Table S1**.

### Bacterial Growth Conditions

Bacteria were grown in LB broth (Lennox) or Mueller Hinton (MH) broth. Antibiotics were used at the following concentrations: carbenicillin (100 μg/mL), chloramphenicol (25 μg/mL for *E. coli*, 5 μg/mL for *V. cholerae*), kanamycin (25 μg/mL), and spectinomycin (50 μg/mL for *E. coli*, 100 μg/mL for *V. cholerae*). Diaminopimelic acid (DAP) was added at 0.3 mM, 5-bromo-4-chloro-3-indolyl-beta-D-galactopyranoside (X-Gal) at 40 µg/mL, glucose at 1% (w/v), arabinose at 0.2% (w/v) and IPTG at 250 μM.

### PCR and Sequencing

For PCRs cloning, Phusion High-Fidelity DNA Polymerase (High-Fidelity Phusion Master Mix, Thermo Fisher #F548S) was used. PCRs were performed on genomic DNA preparations using the DNeasy® Tissue Kit (Qiagen) or on plasmid preparations using the GeneJET Plasmid Miniprep Kit (Thermo Scientific #K0503). PCR products were purified using the GeneJET PCR Purification Kit (Thermo Scientific #K0701).

PCRs for cloning verification were conducted using DreamTaq Polymerase (Thermo Scientific #EP0705). These PCRs were performed directly on bacterial biomass from isolated colonies. Sequencing of PCR products was carried out by Eurofins, using their Sanger sequencing TubeSeq service.

### Plasmid Construction

The plasmids used in this study are listed in **Table S1**. Briefly, plasmids with an oriV(SC101) origin of replication were derived from pMP400, with pBAD promoter and *araC* removed. Plasmids with an oriV(RK2) origin of replication were derived from pSEVA228 (Calero *et al*., 2016), with *xylS* removed. Plasmids for Tn7 transposition were derived from pMP234 (de Lemos Martins *et al*., 2018), and protein expression plasmids were derived from pET22b+ with the *pelB* signal sequence for periplasmic secretion removed (Novagen #69744).

Plasmid constructions were performed using Gibson Assembly (Gibson *et al*., 2009) for complex cloning and inverse PCR followed by ligation for point mutations. Gibson Assembly was carried out as described by (Rabe and Cepko, 2020). Inverse PCR was performed with Phusion High-Fidelity DNA Polymerase (Thermo Fisher #F548S), and the resulting products were ligated using T4 DNA Ligase (Thermo Scientific #EL0011).

Constructed plasmids were transformed into *E. coli* cloning strain NGEpir containing the RK2 plasmid integrated into its genome, allowing the transfer of the generated plasmids via bacterial conjugation. Transformations were performed by electroporation at 1.8 kV/cm, and transformants were selected on MH agar plates supplemented with the appropriate antibiotics and substrate.

For plasmids containing peptide deformylases genes within integron cassettes (pMLB42 to pMLB44), the entire cassette was synthesized as gBlocks Gene Fragments (IDT) and cloned into the pCR-Blunt II-TOPO vector using the Invitrogen Zero Blunt™ TOPO™ PCR Cloning Kit (#450245). These constructs were transformed into *E. coli* cloning strain π3813. Quality controls of all plasmids constructs was done by Sanger sequencing and restriction digestion analysis.

### Bacterial Conjugation

Plasmid transfer was performed by bacterial conjugation using an *E. coli* auxotrophic for diaminopimelic acid (DAP) and containing an integrated RK2 plasmid in the chromosome (*E. coli* NGEpir strain), which allows the mobilization of plasmids with an RK2/RP4 origin of transfer (*oriT*). The donor strain also carries the *pir* gene for the replication of plasmids with an R6K origin.

Conjugation was conducted on Mueller-Hinton (MH) agar plates supplemented with DAP for at least 3 hours. After mating, transconjugants were selected on appropriate antibiotic plates without DAP to eliminate the donor strain.

### Creation of *def1_VCH_* and *def2_VCH_* Deletion Mutants

Deletion mutants of *def1_VCH_* (VC_0046) and *def2_VCH_* (VCA_0150) were generated in *Vibrio cholerae* serotype O1 biotype El Tor strain N16961 *hapR*+ using natural co-transformation, as described by (de Lemos Martins *et al*., 2018). Briefly, *V. cholerae* cells were rendered competent by growth on chitin, following the protocol by (Meibom *et al*., 2005). Once competent, a small amount of PCR product containing an antibiotic resistance cassette flanked by 6 kb of homologous DNA from the intergenic region VC1903/1902, along with an excess of PCR product containing the desired deletion fragment, was added to the medium. This method relies on the principle that if a cell successfully integrates the resistance cassette, it is likely to have also been transformed by the excess deletion fragment (Dalia *et al*., 2014).

To delete *def1_VCH_*, 100ng of a spectinomycin resistance cassette (*aadA7*) was co-transformed with 1µg of a DNA fragment containing the desired deletion. Specifically, the homologous regions consisted of 3 kb upstream of the ATG start codon and 3 kb downstream of the stop codon of *def1_VCH_*, ensuring a total of 6 kb of flanking sequences. Transformants were selected on spectinomycin plates, and correct genome editing was confirmed by PCR and sequencing of the resulting loci. Approximately 50% of the bacteria that integrated the resistance cassette also incorporated the deletion.

The *def2_VCH_* deletion was performed similarly, but with the presence of a thermosensitive pSC101 plasmid expressing *def2_VCH_* under a strong constitutive promoter (pMLB53-TS). The co-transformation and selection were carried out at 30°C and the media supplemented by 50µg/mL of carbenicillin to maintain the expression plasmid. As for *def2_VCH_* deletion mutant, transformants were selected on spectinomycin plates, and correct genome editing was confirmed by PCR and sequencing of the resulting loci. Approximately 50% of the bacteria that integrated the resistance cassette also incorporated the deletion. After *def2_VCH_* was deleted, the thermosensitive plasmid was cured by shifting the temperature from 30°C to 42°C.

For the double deletion mutant (*def1_VCH_* and *def2_VCH_*), *def1_VCH_* was first deleted using the *aadA7* cassette. Following this, *def2_VCH_* was deleted using the strategy described for the single *def2_VCH_* deletion, but this time using a chloramphenicol resistance cassette (*cat*) to replace the spectinomycin resistance cassette at the same integration site. The thermosensitive plasmid pMLB53-TS was kept in the double mutant to ensure the presence of at least one functional PDF, which is essential for bacterial growth, by growing the mutants at permissive temperature.

### Tn7-insertion

For ectopic complementation with PDF genes of the different mutants, we inserted a copy of the gene into the *att-Tn7* site present on the chromosomes of either *E. coli* or *V. cholerae*, near the *glmS* gene following the strategy described in (de Lemos Martins *et al*., 2018). The helper plasmid pMVM1 was first transformed into both *V. cholerae* N16961 and *E. coli* MG1655. This plasmid has a thermo-sensitive pSC101 origin of replication and carries a pBAD promoter that induces the expression of TnsABCD transposases, which catalyze insertion into *att-Tn7* at high frequency as described by (McKenzie and Craig, 2006).

A second shuttle vector derived from pMP234, was used for the specific integration of the different PDF genes (Fournes *et al*., 2021). This vector contains the Tn7 IR sequences recognized by the transposases, and was modified to carry the specific PDF gene along with an *aph* cassette for kanamycin resistance, which allows selection for transposition events. The pMP234 vector is a conditionally replicative vector, which cannot replicate in recipient cells that lack Π protein.

For transposition, the pMP234 shuttle vector was delivered by conjugation into the recipient strains containing pMVM1. Transposition events were selected by plating conjugants on kanamycin plates without DAP. These plates were incubated overnight at 42°C to eliminate the helper vector. All the PDF gene integrated in the chromosome using this set-up are controlled by the same promoter, that of the *V. cholerae def2_VCH_* gene.

### Minimum Inhibitory Concentration (MIC) Determination

For ACT (Sigma-Aldrich #A6671) and LBM415, MICs were determined by broth microdilution according to EUCAST guidelines, as summarized by (Kowalska-Krochmal and Dudek-Wicher, 2021). First ACT and LBM415 were diluted in DMSO to concentrations of 26 mM and 23 mM, respectively. Bacteria overnight cultures were adjusted to an OD600 of 1 and diluted 1:1000 in MH broth (approximately 5×10^5^ CFU/mL). These dilutions were placed in Falcon® 96-well Clear Flat Bottom TC-treated Culture Microplates with Lids (#353072), and the corresponding antibiotic concentrations were added. For control wells without antibiotics, DMSO was added to match the highest volume used in the antibiotic wells to ensure that DMSO itself did not have a toxic effect. The plates were incubated at 37°C with agitation in a TECAN® infinite M200 Pro plate reader for 24 hours and the absorbance at 600 nm was measured every 15 minutes. The MIC was defined as the lowest concentration of antibiotic at which no bacterial growth was observed. Each experiment was performed in biological triplicates with at least three independent repetitions.

### *In vivo* PDF Activity Assay in *V. cholerae*

To assess the *in vivo* activity of peptide deformylase (PDF) in *V. cholerae*, a double mutant strain (Δ*def1_VCH_* Δ*def2_VCH_*) harboring the plasmid pMLB53-TS, which expresses *def2_VCH_* in trans under the expression of a constitutive P_trc_ promoter, was used. A second plasmid with a compatible oriV-trfA origin of replication (backbone pSEVA228) encoding the PDF of interest, under the expression of the constitutive P*_def2VCH_* promoter, was introduced via conjugation. The bacteria containing both plasmids were grown overnight at 30°C with appropriate antibiotics.

Overnight cultures were serially diluted, and 10 µL of each dilution was spotted onto MH agar plates supplemented with kanamycin to maintain the presence of the pSEVA228 plasmid. Plates were incubated at 30°C and 42°C. Growth at 42°C indicated active PDF expression from the pSEVA228 plasmid, as bacteria with inactive PDF would be unable to grow at the higher temperature.

### Computational Genomic Analyses

#### PDF Identification and Annotation Verification

The presence of peptide deformylases (PDFs) was identified in bacterial genomes using annotations from the NCBI Prokaryotic Genome Annotation Process (PGAP). PDFs were extracted using Biopython (v.1.79), and annotation accuracy was confirmed with BLASTp (v.2.16.0) and HMMER (v.3.4) searches (profile PF01327) with the --cut_ga option to ensure high-confidence hits, yielding consistent results across methods.

#### Databases Used

To conduct a comprehensive analysis of peptide deformylases (PDFs) across various bacterial species, multiple genomic databases were employed. For species-wide analysis, reference genomes from RefSeq were utilized, encompassing a total of 5,042 genomes as of May 4, 2024.

To examine intra-species variability, complete genomes available in RefSeq were analyzed, consisting of 40,298 genomes as of April 28, 2024. Plasmid-borne PDFs were identified using the PLSDB, which included 50,554 replicons as of November 23, 2023.

Additionally, the presence of PDFs within integron cassettes was investigated using an in-house database comprising 115,707 proteins. This database was derived from bacterial complete genomes in RefSeq totaling 32,798 genomes as of May 15, 2023, and was analyzed using Integron Finder 2.0.5 (Néron *et al*., 2022) to identify integrons and retrieve cassette information. We used default parameters, with --local-max for accurate detection and --calin-threshold 1 to also detect SALIN (Single *attC* site lacking integron-integrase). For each genome, integrons were detected replicon by replicon. IntegronFinder provides three subtypes of integron-like elements: complete integrons (IntI plus a few cassettes flanked by *attC* sites), CALINs (Clusters of *attC* sites lacking an integrase), and loner integrases. A CALIN with only one *attC* is classified as a SALIN. In the current study, we considered every cassette protein belonging to integrons or CALINs detected by IntegronFinder.

For a detailed analysis of PDFs within the *Vibrionaceae* family, genomes of 136 species with complete sequences available in RefSeq as of January 3, 2023, were examined. The exact list of these genomes is provided in **Table S2**.

#### PDF Genomic Context Analysis

To investigate the genomic context of PDF genes, we extracted the five coding sequences (CDSs) immediately upstream and downstream of each PDF gene across the bacterial reference genomes. Gene annotation was performed using Prokka (v1.14.6).

For the identification of Methionyl-tRNA formyltransferase (*fmt*) genes, we retrieved all CDSs annotated as *fmt* by Prokka that were located directly downstream of PDF genes and encoded on the same DNA strand. These Fmt protein sequences were aligned using MAFFT (--auto), and poorly aligned regions were trimmed with TrimAl. A hidden Markov model (HMM) profile was then constructed and used to search for Fmt homologues using HMMER v3.3.2. Although the HMM profile also detects structurally related formyltransferases (e.g., PurN, PurU, ArnA), applying a stringent e-value cutoff of <10^-20^ excludes these homologues and retains only bona fide Fmt proteins. However, we note that the use of this strict e-value threshold may lead to the omission of highly divergent Fmt sequences, particularly from phylogenetically distant taxa relative to those used to construct the HMM profile.

#### Plasmid Mobility Analysis

Plasmid mobility was assessed using MacSyFinder 2.1.3 with the CONJScan model (Guglielmini *et al*., 2014; Néron *et al*., 2023). Plasmids encoding PDFs were classified into mobility types as follows: (i) conjugative plasmids (pCONJ), which encode a complete mating pair formation (MPF) system along with a relaxase; (ii) decayed conjugative plasmids (pdCONJ), which encode an incomplete MPF system but retain a relaxase; and (iii) mobilizable plasmids (pMOB), which encode a relaxase but lack MPF components. Plasmids carrying only an *oriT* were identified using BLASTn (v2.15.0) as described by (Ares-Arroyo *et al*., 2024). Mobilizable plasmids encoding either a relaxase or an *oriT* can be horizontally transferred by hijacking the conjugation machinery of co-resident conjugative elements.

#### Core Genome Analysis

The core genome of Vibrionaceae was determined using PanACoTA v1.3.1 with a sequence identity threshold of 65% (-i 0.65) and a presence criterion in at least 95% of the analyzed genomes (-t 0.95). The dataset included 136 complete Vibrionaceae genomes (**Table S2**), resulting in the identification of 735 persistent gene families. Each family contained exactly one representative in at least 130 of the 136 genomes (95.0%), while the remaining genomes lacked homologs. Core genome alignment was performed using MAFFT v7.467 with the --auto option. A maximum-likelihood phylogenetic tree was then constructed with IQ-TREE v2.0.6 (Minh *et al*., 2020) using 1,000 ultrafast bootstrap replicates (Hoang *et al*., 2018) and the -m TEST algorithm to determine the best-fit model, which was identified as GTR+F+I+G4 according to the Bayesian Information Criterion (BIC).

#### PDFs Phylogenetic Analysis

For the phylogenetic analysis of peptide deformylases (PDFs), amino acid sequences were similarly aligned using MAFFT (option --auto, v.7.467), or using MUSCLE super5 (v.5.1) to generate Multiple Sequence Alignments (MSAs) by permuting the guide tree (“none”, “abc”, “acb”, “bca”) (Edgar, 2022; Katoh and Standley, 2013). Phylogenetic tree was constructed with IQ-TREE (v.2.2.2.2). This time, 1232 protein models were tested and the best-fitting model was selected according to the BIC. The robustness of the tree was evaluated using 1,000 bootstrap replicates and the SH test replicates.

### Pfold Determination

The folding probability of integron cassette *attC* sites, called Pfold, was determined as described in (Loot *et al*., 2017; Vit *et al*., 2020). Briefly, the Vienna RNA fold WebServers (version 2.6.3) were used with the option “avoid isolated base pair” unchecked, and the DNA parameters model (Matthews model, 2004) selected. Pfold was calculated using the formula: *pFold = eΔGu-ΔGc/RT*. Where ΔGu is the Gibbs energy unconstrainted, ΔGc is the Gibbs energy constrained, R is the gas constant in kcal/mol/K (1.9120458 x 10^-3^), and T is the temperature at 37°C in Kelvin (310.15).

### *attC*-*attI* Recombination Frequency

The *attC-attI* recombination frequency was determined as described in (Vit *et al*., 2020; Vit *et al*., 2021). Briefly, *attC* sites were cloned into a conditionally replicative (suicide) plasmid derived from pSW23T (oriR6K), which codes for the *cat* gene conferring chloramphenicol resistance. This plasmid, which cannot replicate in recipient cells lacking the Π protein, was maintained in the donor strain *E. coli* β2163, which encodes the *pir* gene and can transfer the plasmid via conjugation. Two different *attC* sites were tested: the classical one from the *aadA7* cassette and the one from the integron cassette of *P. suwonensis* 11-1 coding for a PDF. Each site was cloned in both orientations so that either the top or the bottom strand of the attC site is delivered in the recipient strain.

The recipient strain is an *E. coli* MG1655 harboring two plasmids. One plasmid derived from pBAD43, containing the *aadA7* gene and an arabinose-inducible promoter allowing the expression of an integrase *intI1* gene (pL294) or empty (pL290). The second plasmid is derived from pSU38 and contains the *aph* gene and an *attI1* site (p929) allowing recombination with the *attC* site from the suicide plasmid.

The suicide plasmid was transferred by conjugation to the recipient strain on MH agar supplemented with DAP and arabinose allowing the expression of the integrase. The suicide plasmid cannot replicate in the recipient strain, so maintaining chloramphenicol resistance requires recombination between the *attC* site on the suicide plasmid and the *attI* site on plasmid p929, forming a chimeric plasmid. In the absence of integrase, this recombination is not possible.

Following conjugation, transconjugants were selected on MH agar supplemented with chloramphenicol, kanamycin, and spectinomycin. Recipient cells were selected on MH agar with kanamycin and spectinomycin. 16 transconjugants were tested by PCR to verify the correct *attC*-*attI* recombination using the primers SwBeg-MFD when the *attC* bottom strand is delivered and the primers SwBeg-MRV when the top strand is delivered.

Recombination frequencies were calculated by dividing the number of transconjugants by the total number of recipients. If some of the 16 PCR-tested transconjugants did not show *attC-attI* recombination, the frequencies were recalculated as described by (Vit *et al*., 2020).

### Protein Purification

Four distinct plasmids were constructed for the expression of peptide deformylases, each featuring an N-terminal hexahistidine tag (6xHis) followed by a 3xGS linker. The plasmids and their encoded proteins are detailed in supplementary **table S1**.

#### Protein expression

*E. coli* Bli5 (BL21 (DE3) + pDIA17) (Munier *et al*., 1992) cells transformed with the respective pET22b plasmid encoding peptide deformylase (pMLB109, pMLB110) were grown in 1L of LB medium supplemented with chloramphenicol and ampicillin at 37°C in a 5L Erlenmeyer flask until the optical density at 600 nm (OD600) reached 0.8. Protein expression was induced by adding IPTG to a final concentration of 250 µM, and the culture was incubated for an additional 3 hours at 25°C. The cells were harvested by centrifugation at 8000xg for 15 minutes at 4°C, and the resulting cell pellet was stored at −20°C for up to one month prior to lysis.

#### Cell Lysis

The frozen cell pellet was resuspended in 50 mL of lysis buffer (25 mM Tris-HCl, pH 7.8, and 100 mM NaCl) and lysed using a CellD cell disruptor at 1.4 Kbar. The lysate was clarified by centrifugation at 25,000xg for 30 minutes at 4°C.

#### Affinity Chromatography

The clarified lysate was loaded onto a HisTrap™ High Performance 5mL column (Cytiva 17-5248-02) using an Äkta Start system. The column was equilibrated with lysis buffer (25 mM Tris-HCl, pH 7.8, and 100 mM NaCl) before sample application. Following sample application, the column was washed with 10 column volumes of the same buffer at a flow rate of 2 mL/min. Protein elution was carried out with a gradient of buffer A (25 mM Tris-HCl, pH 7.8, 100 mM NaCl, and 20 mM imidazole) and buffer B (25 mM Tris-HCl, pH 7.8, 100 mM NaCl, 300 mM imidazole, and 10% glycerol). The gradient was run over 40 mL, increasing buffer B from 0% to 100% at a flow rate of 2 mL/min, and fractions were collected every 2 mL. The protein of interest started to elute at approximately 50% buffer B (150 mM imidazole) and was concentrated on Vivaspin® 6 Centrifugal Concentrator Polyethersulfone with a Molecular weight cutoff of 10 KDa to a final volume of 5mL.

#### Size Exclusion Chromatography

For further purification, the 5 mL protein solution from affinity column chromatography was subjected to size exclusion chromatography using a HiLoad 16/600 Superdex 75 pg column on an UPC-900 system. The column was equilibrated with lysis buffer (25 mM Tris-HCl, pH 7.8, and 100 mM NaCl), and the protein was eluted in the same buffer at a flow rate of 1mL/min. Fractions containing the target protein were collected, pooled, and concentrated on Vivaspin® 6 Centrifugal Concentrator Polyethersulfone with a Molecular weight cutoff of 10 KDa to a final concentration of 10mg/mL for downstream applications.

#### SDS-PAGE Analysis

Fractions were analyzed by SDS-PAGE using Mini-PROTEAN TGX Stain-Free gels (4-15% acrylamide, Bio-Rad) in a running buffer of 1X Glycine-Tris-SDS (19.2 mM glycine, 2.5 mM Tris, and 0.01% SDS, pH 8.3). Electrophoresis was performed at 170 V, and the gel was stained with Coomassie Brilliant Blue R-250 Staining Solution (Bio-Rad #1610436) for 20 min followed by destaining with Coomassie Brilliant Blue R-250 Destaining Solution (Bio-Rad #1610438) to assess protein purity.

#### Quality control Analysis

The quality control of the recombinant proteins was assessed by mass spectrometry and dynamic light scattering (DLS) according to the ARBRE-MOBIEU/P4EU guidelines (de Marco *et al*., 2021).

### Crystallization

Screening of crystallization conditions and optimization of hits were performed at the Crystallography Core Facility of the Institut Pasteur (Weber *et al*., 2019). Briefly, initial screenings were performed in 96-well Greiner plates using a Mosquito automated nanoliter dispensing system (TTP Labtech, Melbourn, UK). The plates were then stored at 18 °C in a RockImager automated imaging system (Formulatrix, Bedford, USA) to monitor crystal growth. Initial crystallization hits were optimized in 24-well plates using the hanging drop method. The best Def1_VCH_ crystals grew in wells containing 20% (w/v) PEG4K, 20% (v/v) 2-Propanol and 0,1M Na3-citrate pH 5.6 in the reservoir. Def2_VCH_ crystals were obtained in 20% (w/v) PEG4K, 10% (v/v) 2-Propanol, 0,1M Hepes pH 7.5. The crystals were flash cooled in liquid nitrogen for data collection using the crystallization solution as cryoprotectant for Def1_VCH_ and a mixture of oils (50% paratone N + 50% paraffin) for Def2_VCH_.

### Diffraction data collection and structure determination

X-ray diffraction data were collected at beamlines PROXIMA 1 and PROXIMA 2a (Synchrotron SOLEIL, St. Aubin, France) and processed with autoPROC (Vonrhein *et al*., 2011). The crystal structures were solved by the molecular replacement (MR) method using Phaser (McCoy *et al*., 2007), and trimmed AlphaFold (Jumper *et al*., 2021; Mirdita *et al*., 2022) models as search probe. The final models were obtained through interactive cycles of manual model building with Coot (Emsley and Cowtan, 2004) and reciprocal space refinement with Buster (Bricogne *et al*., 2017) and Phenix (Liebschner *et al*., 2019). Figures were generated using ChimeraX.

### Accession codes

Atomic coordinates and structure factors have been deposited in the RCSB Protein Data Bank under the accession codes 9QFR (Def1_VCH_-ACT) and 9QFT (Def2_VCH_-ACT).

## Results

### Half of bacterial species harbors several PDF

Despite longstanding knowledge of PDF duplication, no large-scale study has explored yet the distribution of PDFs across bacterial genomes. We analyzed 5,042 fully sequenced genome of bacterial references species available in the RefSeq database as of May 4, 2024 and measured the occurrence of genes coding for PDFs in their genomes (**Fig. 1A**). PDF genes were extracted using annotations from the NCBI Prokaryotic Genome Annotation Process (PGAP). Annotation accuracy was confirmed by BLASTp and HMMER searches (profile PF01327), leading to similar results.

**Fig. 1:**
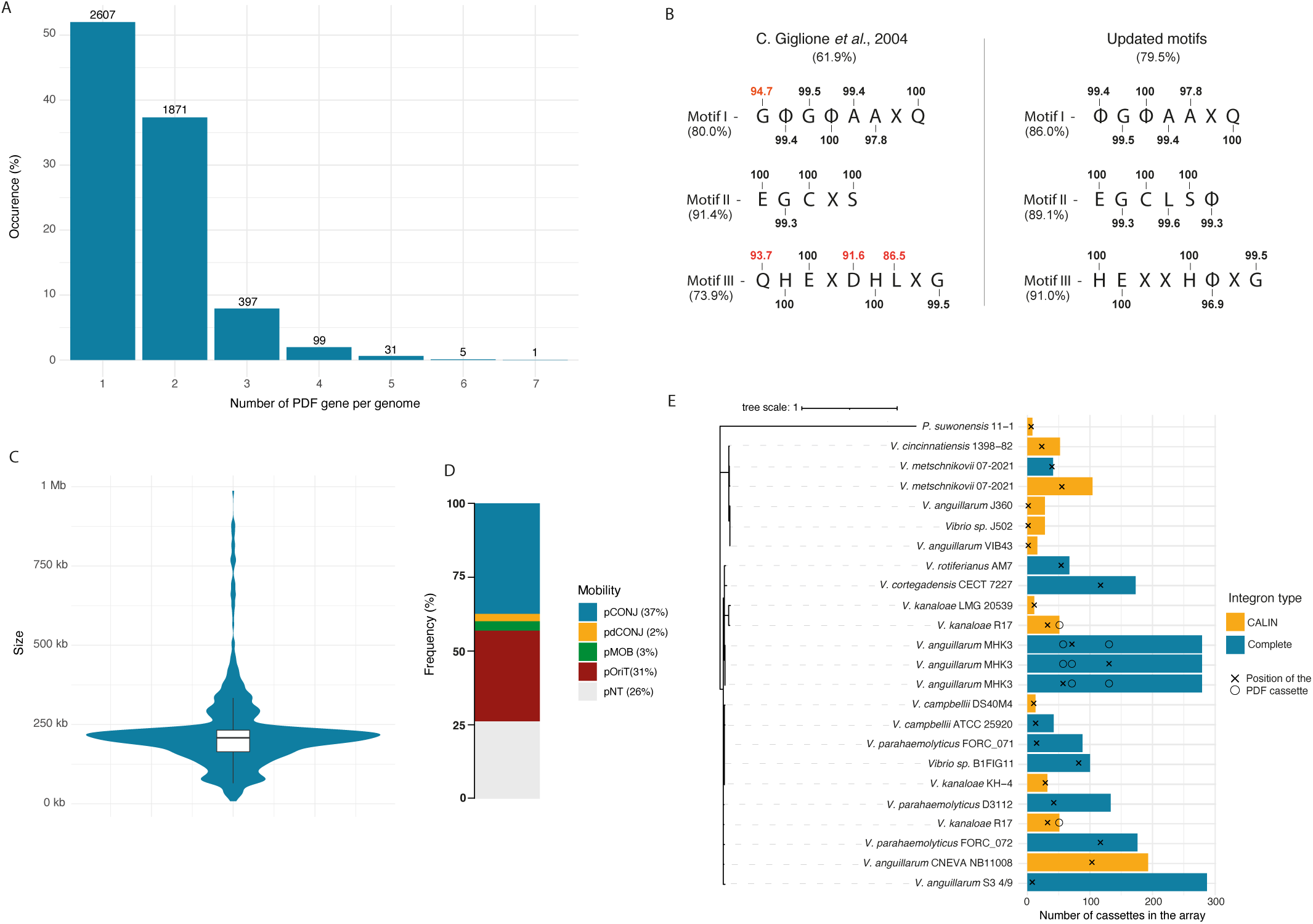
Prevalence and distribution of accessory peptide deformylases in bacterial genomes. **A)** Distribution of Peptide deformylases in bacterial reference genomes. The number of species is indicated above the histograms. **B)** Amino acid conservation is shown for the three motifs originally described by (C. Giglione *et al.,* 2004) and for revised motifs derived from PDFs found in bacterial reference genomes encoding a single PDF copy. The overall conservation of each motif across all PDFs is indicated below, along with the proportion of PDFs that exactly match the canonical or updated motif sequences. Conservation values for individual amino acid positions are indicated above or below the corresponding residue, based on the subset of genomes encoding a single PDF. Amino acid conservation below 95% are highlighted in red. Φ represents a hydrophobic amino acid and X represents any amino acid. **C)** Violin plot illustrating the size distribution of plasmids encoding PDFs. The violin plot shows the density of plasmid sizes, with the white box inside representing the interquartile range (IQR) and the central line indicating the median size. **D)** Bar chart depicting the classification of PDF-encoding plasmids based on their mobility: 36% encode all the key genes required for conjugation, i.e mating pair formation (MFP) machinery and a relaxase (conjugative plasmids: pCONJ), 2% encode an incomplete MFP machinery and a relaxase (decayed conjugative plasmids: pdCONJ), 3% encode only a relaxase (mobilizable plasmids: pMOB), and 31% encode an *oriT* without relaxase (pOriT). **E)** Phylogeny and genomic context of PDFs encoded within integron cassettes. The phylogenetic tree (left) represents the evolutionary relationships among PDF protein sequences encoded within integron cassettes. The bar plot (right) represents the total number of cassettes in each integron, with colors distinguishing complete integrons (blue) from CALINs (yellow). The relative position of the PDF cassette within each integron is indicated by a cross when it corresponds to the sequence used in the phylogeny and by a circle when additional PDF cassettes are present in the same integron. The phylogenetic tree was constructed using IQ-TREE multi-core v.2.2.2.2 (Minh et al., 2020) based on a MAFFT-aligned dataset (Katoh and Standley, 2013). Among 1232 proteins models tested, the Q.plant+G4 model was selected as the best-fitting model according to the Bayesian information criterion using the IQ-TREE-M TESTNEW algorithm. Statistical support was assessed using 1,000 ultrafast bootstrap replicates and 1,000 SH approximate likelihood ratio test replicates (Hoang et al., 2018).

This analysis revealed that 48% of bacterial species harbor more than one PDF. Notably, over 2.7% of reference genomes contain more than three PDF genes, with *Dactylosporangium vinaceum* holding the record with seven PDFs. Among the 5,042 reference genomes, only 31 genomes lacked identifiable PDF genes. Of these, 29 are endosymbiotic bacteria, where the absence of essential genes has been previously documented (McCutcheon and Moran, 2011) and 2 are probably incomplete as they also lack the formyltransferase *fmt* gene thought essential and conserved among all bacteria species (**Table S3**).

### Conservation of essential residues: refinement of the catalytic motifs

The PDF sequence contains three conserved motifs reported to be essential for enzymatic activity: motif I (GΦGΦΑAXQ), motif II (EGCXS), and motif III (QHEXDHLXG), where Φ represents a hydrophobic amino acid and X represents any amino acid (Giglione *et al*., 2004).

To evaluate the potential activity of these PDFs, we analyzed the presence of these three conserved motifs (Giglione *et al*., 2004). Surprisingly, only 61.9% of the PDFs exhibited exact conservation of these motifs. This may suggest that many accessory PDFs might be weakly active or inactive, similar to those identified in *S. aureus* and *S. pneumoniae* and belonging to the type 3 PDF (Margolis *et al*., 2000; Margolis *et al*., 2001; Grzela *et al*., 2017). However, 24.5% of the reference genomes did not encode any PDF matching the exact sequence of these motifs. Given that at least one active PDF is essential for bacterial survival, this suggested that the motifs described by Giglione and colleagues, may not be reliable enough as indicators of PDF activity across all bacterial species. Aligning PDFs from genomes harboring only one PDF revealed that the motifs described by C. Giglione *et al*. (Giglione *et al*., 2004) are indeed not strictly conserved (**Fig. 1B**).

Based on these observations, we propose an update to the motifs described by C. Giglione and colleagues, with a cut-off for the amino acids conserved in more than 95% of PDFs present in genomes with only a single PDF copy: motif I (ΦGΦΑAXQ), motif II (EGCLSΦ), and motif III (HEXXHΦXG). The vast majority of PDFs in bacterial genomes (80%) possess these updated motifs, encompassing the amino acids critical for enzymatic activity (**Fig. 1B**). However, 7.1% of bacterial reference genomes still do not encode a PDF with the exact sequence of these updated motifs indicating that the exact conservation of these three motifs is not essential for the PDF enzymatic activity, or that they contain sequencing errors.

### Conservation and Genomic Organization of Formylation and Deformylation Genes in Bacteria

Although it is widely assumed that the formylation of initiator tRNA^Met^ by Methionyl-tRNA formyltransferase (Fmt) is a conserved feature of bacterial translation, to our knowledge, no recent large-scale study has systematically demonstrated this conservation. Similar to our analysis of PDFs, we screened 5,042 bacterial reference genomes and found that the vast majority (98.6%) encode a single copy of the *fmt* gene. Only ten genomes encode two *fmt* homologs, while the few genomes lacking *fmt* (73 in total) predominantly correspond to obligate endosymbionts with highly reduced genomes.

It is also generally accepted that *fmt* and *def* genes are co-localized and often organized into a single operon, as exemplified in *E. coli* (Meinnel and Blanquet, 1993; Mazel *et al*., 1994; Mazel *et al*., 1997). Genomic context analysis of PDF genes revealed that, although *fmt* and *def* are frequently co-localized, this association is conserved in only 58% of bacterial species. Furthermore, in *E. coli*, the *fmt*-PDF operon additionally includes *rsmB* and *trkA*, encoding proteins involved in 16S rRNA methylation and potassium transport regulation, respectively. This extended operonic structure is restricted to Gammaproteobacteria and found in only 62% of references species within this clade.

Additionally, we identified 145 bacterial species in which PDF genes are located in close proximity to other PDF genes (less than 5 genes apart), consistently on the same DNA strand, suggesting potential co-transcription. Notably, 104 of the 145 species exhibiting clustered PDF genes were found to belong to the order Rhodobacterales. This genomic organization appears to be a conserved feature of this clade, as 98.1% of the Rhodobacterales reference species in our dataset harbor multiple PDF genes arranged in a cluster. In the majority of these Rhodobacterales genomes (62.5%), three PDF genes were found grouped together. Although highly conserved in *Rhodobacterales*, the biological rationale behind the clustering of PDF genes remains elusive.

An unusual case was observed in the four reference genomes of *Exiguobacterium*, a genus from the order *Bacilliales*, in which *def* and *fmt* appear to be fused into a single gene. Interestingly, all *Exiguobacterium* genomes also encode a second PDF containing all residues critical for catalytic activity. Whether this *def-fmt* fusion reflects a true biological feature or results from sequencing or annotation artifacts remains unclear, as does the functionality of the corresponding proteins.

### PDFs are encoded on mobile genetic elements

The analysis of all complete bacterial genomes in RefSeq as of April 28, 2024 (40,298 genomes) distributed among 5,728 species revealed intra-species variability in the number of PDFs encoded within bacterial genomes. Indeed, approximately 10% of bacterial species with at least two complete genomes in this database show no conservation in PDF copy number, suggesting that some PDFs may be associated with mobile genetic elements (MGEs). To further investigate this possibility, we analyzed a plasmid database (PLSDB) and an integron cassette database to determine whether PDFs are encoded within these MGEs.

#### Plasmid

In the plasmid database, we identified 676 replicons harboring at least one PDF gene. However, this database also mis-includes some secondary chromosomes or chromids. Generally, chromosomes and chromids are distinguished from plasmids by their “essential” nature and larger size (Harrison *et al*., 2010; diCenzo and Finan, 2017). Applying an additional filter to remove replicons larger than 1 Mb, we identified 657 replicons smaller than 1 Mb, carrying one PDF genes and 10 harboring 2 PDFs genes (**Table S4**). Analysis revealed that the median size of plasmids encoding a PDF gene is 210 kb (**Fig. 1C**). They are mainly found in *K. pneumoniae* (57.2%) and *E. coli* (25.9%), but they can be found among 58 different bacterial species from 35 genera (**Fig. S1A** and **Table S4**). A significant proportion (67.5%) of the identified plasmidic PDFs are identical to the chromosomal *E. coli* PDF, typically found in an operon with *fmt*, and are distributed across 12 Enterobacteriaceae species spanning the genera *Klebsiella, Escherichia, Shigella, Salmonella,* and *Enterobacter*. This widespread distribution likely results from horizontal transfer, as plasmids can disseminate via conjugation or mobilization. Consistently, approximately 74% of PDF-encoding plasmids were predicted to be mobile (**Fig. 1C-D**). Additionally, at least 5 plasmids have been already identified as phage-plasmids, which are also capable of horizontal transfer between bacteria (Pfeifer *et al*., 2022; Ares-Arroyo *et al*., 2024; Pfeifer and Rocha, 2024).

#### Integron

The integron system is a powerful genetic mechanism that enables bacteria to rapidly adapt to changing environments by capturing, stockpiling, and rearranging gene-encoding cassettes (Escudero *et al*., 2015). These cassettes often contain genes crucial for bacterial adaptation, such as those conferring antibiotic resistance or phage-defense (Darracq *et al*., 2025; Kieffer *et al*., 2025). A promoter, known as the Pc promoter, is located at the beginning of the integron. Typically, cassettes are promoter-less, and their expression relies on their position within the integron. Thus, cassettes positioned near the beginning (*attI* site) are actively expressed, establishing a gradient of expression that decreases with increasing distance from the Pc promoter. Cassettes located far from the promoter are minimally expressed and essentially serve as a low-cost molecular reservoir of functions.

We probed our in-house integron database (see materials and methods) and identified 24 PDF cassettes across 20 bacterial strains, predominantly within the genus *Vibrio* (**Fig. S1B**) (**Table S5**). Notably, two integrons contain multiple PDF cassettes: *V. anguillarum* MHK3 harbors three PDF cassettes and *V. kanaloae* R17 harbors two. In *V. metschnikovii* 07_2421, two cassettes array were found, each containing a PDF cassette. The phylogenetic analysis of integron-encoded PDF showed that all *Vibrio* PDF cassettes share a common origin, likely driven by horizontal transfer. In contrast, the PDF cassette identified in *P. suwonensis* is phylogenetically distinct, and sequence analysis revealed that its closest homolog is a PDF from an Alphaproteobacteria of the genus *Chelatococcus*, with which it shares 51% identity, supporting a separate evolutionary origin (**Fig. 1E**).

PDF cassettes are present in both complete integrons and CALIN (Cluster of *attC* site lacking integron integrase (Cury *et al*., 2016)), but all these strains have an integrase capable of mobilizing cassettes from CALINs. The short distance between the recombination site and the start codon (25-28 bp) likely precludes the presence of an internal promoter within the cassette, making PDF expression likely dependent on its position in the integron. Their positions in the integrons vary, with some located far from the integration site (*attI*), implying ancient acquisition, while others are close to the integration site, suggesting recent acquisition or remobilization, as is the case for the *V. anguillarum* str. S3-4/9 PDF cassette (8th position) (**Fig. 1E**). Additionally, in the CALINs of *Vibrio sp*. J502 and *V. anguillarum* VIB4 and J360, PDF cassettes were found in the first position, downstream of a putative promoter sequence which could lead to their expression as already described by Loot and colleagues (Loot *et al*., 2024).

Most PDF cassettes are located in sedentary chromosomal integrons (SCIs). However, one cassette is found in a mobile integron on a plasmid, in *V. campbellii* DS40M4 and the CALIN identified in *P. suwonensis* resembles a mobile integron that has been integrated in the chromosome, as it is characterized by heterogeneous *attC* sites (60 to 112 bp) and by the presence of the aminoglycoside resistance gene *aadA1*, typically found in mobile integrons, in its first position.

In summary, our results highlight the widespread presence of multiple PDFs in bacterial genomes. Most of these PDFs contain the essential catalytic amino acids, suggesting that they are likely active. In addition, the observation that PDFs are frequently associated with mobile genetic elements capable of horizontal dissemination, reinforce the fact that they likely play an adaptive role.

To further investigate the possible adaptive function of accessory PDFs, we focused on *Vibrio cholerae*, a well-studied pathogen and model organism known to possess two PDFs (Margolis *et al*., 2000). Furthermore, given that integron cassettes carrying PDF are predominantly found in *Vibrio* species, *V. cholerae* offers an ideal system to explore the adaptive roles of accessory PDFs.

### *Vibrionaceae* PDF diversity

We investigated the diversity of PDFs in Vibrionaceae species by analyzing a subset of 136 complete genomes available in RefSeq (**Table S2**). Among these, 46.7% possess multiple PDF genes, with 12 species, primarily from the *Photobacterium* genus, containing three PDFs (8.8%) (**Fig. S2**).

Phylogenetic analysis of the PDFs from these 136 genomes, along with the 23 integron-associated PDFs that were not found in these reference genomes, revealed several distinct subfamilies of proteins (**Fig. 2**). All species share a conserved canonical type 1b PDF, designated as Def1, derived from a common ancestor and found in an operon with the formyltransferase Fmt, as previously observed in other bacteria (Meinnel and Blanquet, 1993; Mazel *et al*., 1997).

**Fig. 2:**
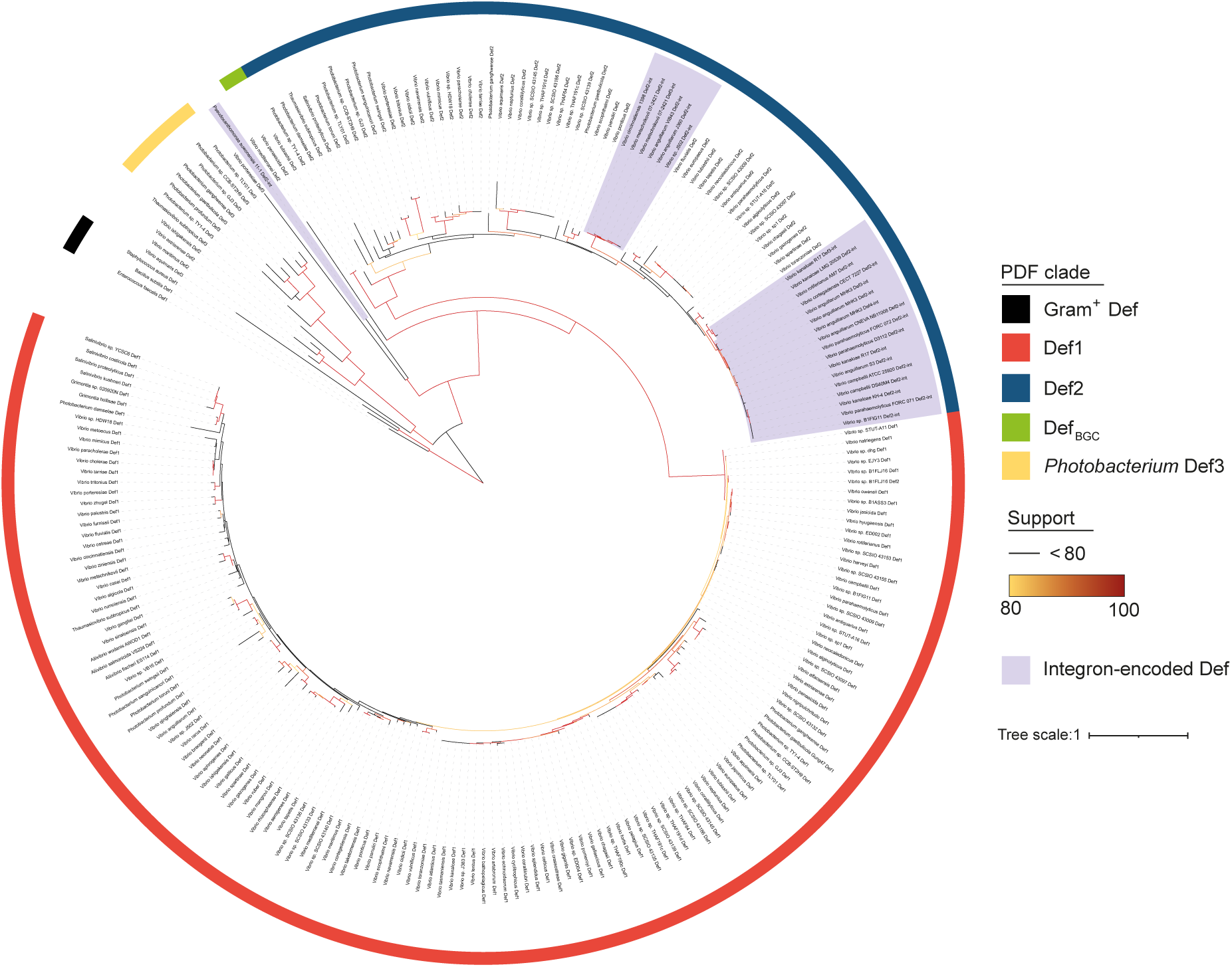
Phylogenetic tree of integron-encoded and *Vibrionaceae* PDF. The phylogenetic tree represents PDFs from 136 *Vibrionaceae* genomes (listed in **Table S2**) and 24 PDFs from integron cassettes (listed in **Table S5**). The colored outer ring indicates the PDF clades. UFBoot values over 80 are shown in a yellow-red gradient. The Tree was rooted using tree Def sequences from Gram positive bacteria as an outgroup.

The second major clade, designated as Def2, includes PDFs found in *Vibrio* integron cassettes as well as most accessory PDFs identified in Vibrionaceae, including the type 1b accessory PDF previously described in *V. cholerae,* named Def2_VCH_ (Margolis *et al*., 2000). This group, present in 41.6% of Vibrionaceae (**Fig. S2**), exhibits greater sequence variability than Def1, suggesting a possible adaptive role. Motif conservation analysis revealed that 61.8% of Def2 variants carry a mutation in motif I, where the second alanine is replaced by a serine— a modification that does not affect enzymatic activity (see below).

Given the widespread distribution of Def2 PDFs across the Vibrionaceae family, we conducted a phylogenetic analysis to determine whether the phylogenetic tree of the Def2 proteins matches the phylogeny of the species in which the proteins are found. Our analysis showed that the phylogenetic distribution of Def2 PDFs is not congruent with the species phylogeny, supporting the idea that the accessory Def2 PDF was acquired via horizontal gene transfer between the ‘cholerae’ and ‘vulnificus’ strains (**Fig S2 and S3**).

We also found that the previously identified PDFs in *V. tubiashii* and *V. penaeicida*, situated within a biosynthetic gene cluster (BGC) that includes two non-ribosomal peptide synthetases (NRPS), a type 1 polyketide synthase (PKS), and a methyltransferase (Cordero *et al*., 2012; Costa *et al*., 2024) are closely related to the Def2 group (**Fig. 2**). It has been suggested that this BGC might be involved in synthesizing a peptide deformylase inhibitor, with the BGC-encoded PDF serving as an antidote (Costa *et al*., 2024). Recent functional studies have confirmed this hypothesis, showing that the encoded metabolite indeed inhibits PDF by binding its active site (Chen *et al.,* 2025; Rill *et al*., 2025). Furthermore, inhibition of PDF was found to promote the activation of specific prophages in *Vibrio*, through a mechanism that bypasses the canonical SOS pathway (Chen *et al.,* 2025).

Unfortunately, Costa and colleagues were not able to validate this hypothesis. However, emerging data strongly suggest that this cluster indeed produces a PDF inhibitor. Beyond these major PDF clades, we identified additional PDFs that are phylogenetically distinct from both the canonical Def1 and the accessory Def2 clades (**Fig. 2**). Since *Vibrio* integron PDFs belong to the Def2-like group, we focused on characterizing the role of accessory Def2 PDFs in *V. cholerae*.

### *Vibrio cholerae* harbor 2 actives PDF

*V. cholerae* possesses two PDFs, Def1_VCH_ and Def2_VCH_. The canonical PDF, Def1_VCH_, is located on chromosome 1 within an operon that includes *fmt*, *rsmB*, *trkA* (as in *E. coli* (Cho *et al*., 2009)), *trkH*, and a gene of unknown function with a conserved DUF3157 domain (VC0041). The *rsmB* gene encodes a methyltransferase that specifically methylates cytosine 967 (^m5^C967) of the 16S rRNA, stabilizing the binding of fMet-tRNA^fMet^ to the 30S pre-initiation complex before start codon recognition (Burakovsky *et al*., 2012). The *trkA* and *trkH* genes are involved in potassium transport regulation (Zhang *et al*., 2020). In contrast, the accessory PDF, Def2_VCH_, is encoded on chromosome 2 as a monocistronic unit. According to previously published proteomic data from the *Vibrio cholerae* N16961 WT strain grown in MH medium at 37°C, the PDF Def1_VCH_ is 2 times more abundant than Def2_VCH_ (Fruchard *et al*., 2025).

Despite sharing only 51% amino acids sequence identity, Def1_VCH_ and Def2_VCH_ exhibit significant structural homology, with their most notable differences located in their C-terminal regions (**Fig. S4A**). Amino acid sequence alignment reveals strict conservation of the three motifs that have been described as critical for catalytic activity. However, in Def2_VCH_, shows a specific pattern variations in the amino acids, described in *E. coli*, as responsible for ribosome binding (**Fig.S4B**). Specifically, the C-terminal helix of the *E. coli* PDF has been shown to insert into a groove between ribosomal proteins uL22 and uL32 (Bingel-Erlenmeyer *et al*., 2008; Sandikci *et al*., 2013; Akbar *et al*., 2021). The first half of this helix interacts with uL22 through residues L149, K150, R153, and K157, while the second half interacts with 23S rRNA through K160 and R163 (Bingel-Erlenmeyer *et al*., 2008; Akbar *et al*., 2021). In Def1_VCH_, all amino acids crucial for ribosome binding are identical to *E. coli* PDF, while in Def2_VCH_, R153 is substituted by a methionine and the R163 by a lysine (**Fig. S4B**). In *E. coli*, R153 forms a hydrogen bond with E52 of ribosomal protein uL22, and a methionine substitution likely disrupts this interaction. Interestingly, all PDFs within the Def2 group lack R153 (**Fig. S4C**), instead they have a methionine in 95% of cases, and more rarely an isoleucine or leucine, all hydrophobic amino acids (**Fig. S4D**). Meanwhile, the glutamate E52 is conserved in 95% of *Vibrionaceae* uL22 proteins, the others have an aspartate (4%) or valine (1%) as this position. This suggests a possible functional divergence in their interaction with the ribosome.

We successfully deleted either *def1_VCH_* or *def2_VCH_*, suggesting that both PDFs are active. To further assess their essentiality, we attempted to construct a double mutant lacking both genes. Since PDF activity is known to be essential in many bacteria, we hypothesized that such a mutant would not be viable. To circumvent this, we maintained deformylase activity *in trans* using a thermosensitive plasmid to construct the double deletion mutant. To assess whether each PDF is functional, we introduced a compatible plasmid expressing either *def1_VCH_* or *def2_VCH_* and shifted the temperature to 42°C to cure the thermosensitive plasmid. Growth at 42°C was only observed when the second plasmid carried an active PDF, confirming that both Def1*_VCH_* and Def2*_VCH_* are enzymatically functional and can individually support bacterial viability (**Fig. 3A**).

**Fig. 3:**
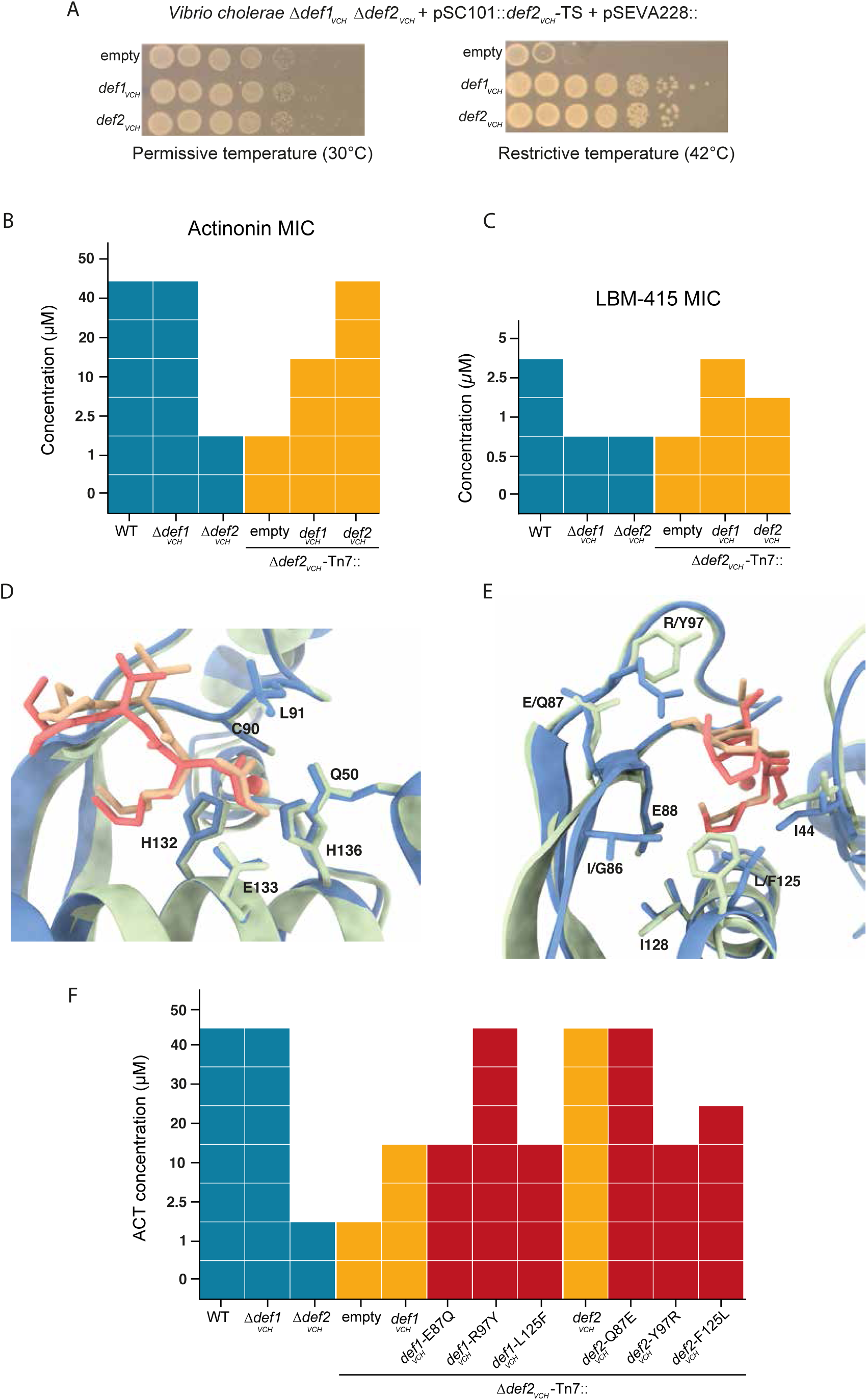
Def2 confers resistance to ACT. **A)** *In vivo* PDF activity assay. Drop test of *V. cholerae* Δ*def1_VCH_* Δ*def2_VCH_* harboring a pSC101::*def2_VCH_*-TS plasmid and a pSEVA228 plasmid at permissive temperature (left panel) or restrictive temperature (right panel). Growth at restrictive temperature indicate that the PDF coding in the pSEVA228 plasmid is enzymatically active. **B)** Antibiotic susceptibility matrix of ACT. CMI was determine by broth microdilution as the minimal concentration of ACT with no observed growth. **C)** Antibiotic susceptibility matrix of LBM415. CMI was determine by broth microdilution as the minimal concentration of LBM415 with no observed growth. **D-E)** Structural superposition of Def1_VCH_ (blue) and Def2_VCH_ (green) in complex to ACT (Def1_VCH_ -ACT: red, Def2_VCH_ -ACT: orange). Side chains of amino acids involved in ACT and/or metal cofactor binding are shown in sticks representation. **F)** Impact of the amino acid substitution on the resistance level to ACT. Antibiotic susceptibility matrix of ACT. CMI was determine by broth microdilution as the minimal concentration of ACT with no observed growth.

### Def2 reduce susceptibility to ACT

Actinonin, the first discovered PDF inhibitor, is a natural antibiotic synthesized by a BGC consisting of non-ribosomal peptide synthetases (NRPS) and polyketide synthases (PKS), identified in *Streptomyces* species. It is plausible that ACT, along with other yet-to-be-identified natural PDF inhibitors, is present in the environment, where it may influence microbial competition. Integrons, which play a pivotal role in bacterial adaptation to environmental pressures, frequently harbor genes linked to antibiotic resistance and phage-defense (Darracq *et al*., 2025; Kieffer *et al*., 2025). Homologs of Def2_VCH_ are found within NRPS and PKS operons in some *Vibrio* species (**Fig. 2**) and in both chromosomal and mobile integrons (**Fig. 2**). These observations led us to hypothesize that accessory PDFs, such as Def2_VCH_, might contribute to resistance against PDF inhibitors.

To test this hypothesis, we investigated whether the presence of the accessory PDF Def2_VCH_ provides an advantage in resistance to ACT. Deletion of the canonical PDF *def1_VCH_* did not impacted the sensitivity to ACT, but the deletion of the accessory PDF *def2_VCH_* lead to a 20-fold decrease in the MIC of ACT. Ectopic complementation of the Δ*def2_VCH_* strain with a *def2_VCH_* gene integrated in the chromosome restored sensitivity to levels observed in the wild-type (WT) and Δ*def1_VCH_* strains, while complementation with the *def1_VCH_* gene resulted in a more moderate gain in resistance (**Fig. 3B**). We further assessed resistance to another peptidomimetic PDF inhibitor, LBM415, which is derived from ACT and engineered by Novartis (**Fig. S5**) (Anderegg *et al*., 2003; Fritsche *et al*., 2005; Rolan *et al*., 2011). In contrast, deletion of either *def1_VCH_* or *def2_VCH_* led to a comparable decrease in LBM415 resistance. Complementation of the Δ *def2_VCH_* mutant with *def1_VCH_* conferred only a slight increase in resistance compared to complementation with *def2_VCH_*, suggesting a more limited protective effect of the canonical PDF against this inhibitor. (**Fig. 3C**).

Given the observed difference in resistance, we hypothesized that Def2VCH may exhibit altered binding affinity for ACT compared to Def1VCH. To explore this possibility at the molecular level, we determined high-resolution crystal structures of both proteins in complex with ACT (**Fig. 3D-E** and **Table S6**). In both structures, ACT binds in the same location within the catalytic pocket and adopts a nearly identical orientation and conformation, blocking access to the active site located at the bottom of a deep cavity in the protein core (**Fig. S6A-B**).

In Def1_VCH_, all previously identified amino acids involved in ACT binding in *E. coli* PDF were also identified interacting with ACT (**Fig. 3D-E** and **Fig. S4**) (Clements *et al*., 2001). In contrast, Def2_VCH_ exhibited a different binding pattern. This is mainly due to the substitution of arginine 97 with a tyrosine, which weakens the interaction with ACT by abolishing important hydrogen bonds between the NH1 and NH2 atoms of Arg97 and solvent exposed oxygen atoms of ACT (namely O20 and O27). Additionally, in the hydrophobic pocket containing the ACT P1’ aliphatic chain, isoleucine at position 86 and leucine at position 125 in Def1_VCH_ are substituted by glycine and phenylalanine, respectively, in Def2_VCH_ (**Fig. 3D-E** and **Fig. S4B**). These substitutions likely contribute to the variation in ACT binding affinity between Def1_VCH_ and Def2_VCH_, explaining their different capacities to confer resistance to this inhibitor.

To further investigate the role of these amino acid substitutions in ACT resistance, we engineered reciprocal mutations, swapping the residues implicated in ACT interaction between Def1_VCH_ and Def2_VCH_. Substitution of arginine 97 with tyrosine in Def1_VCH_ resulted in a significant increase in the MIC to ACT, reaching levels comparable to those conferred by Def2_VCH_ (**Fig. 3F**). Conversely, substitution of tyrosine 97 with arginine in Def2_VCH_ led to a marked reduction in the MIC, aligning with the resistance phenotype observed for Def1_VCH_ (**Fig. 3F**). Substitutions of glutamate 87 to glutamine (E87Q) and leucine 125 to phenylalanine (L125F) in Def1_VCH_ had either no or intermediate effects, in the same direction as the R97Y substitution, and their effects appeared to be cumulative (**Fig. 3F**). Those substitutions are conserved among the whole Def2 group and among BGC-encoded PDF identified in *Vibrio* species. It is therefore likely that theses accessories PDFs, including the integron-encoded one, may also confer a better resistance to actinonin, or other PDF inhibitors, in their native host.

These results indicate that the accessory PDF, Def2_VCH_, plays a crucial role in resistance to actinonin in *V. cholerae*. The resistance conferred by Def2_VCH_ appears to be linked to a lower affinity of ACT for this protein compared to the canonical PDF, Def1_VCH_. ACT is a natural product produced by a BGC in *Streptomyces* bacteria, suggesting that PDF inhibitors may be present in the environment and play a role in bacterial competition.

### Integron PDF are enzymatically active

We tested the enzymatic activity of three PDF cassettes: VaDef from *V. anguillarum* str. MHK3, VkDef from *V. kanaloae* str. R17, and PsDef from the integron of *Pseudoxanthomonas suwonensis* str. 11-1. This was done *in vivo* using the previously described V*. cholerae* mutant lacking both PDF genes. All three were found to be functional and able of supporting viability in *V. cholerae* (**Fig 4A**). As all integron-encoded Def2-like proteins exhibited a substitution of the second alanine (A48) in motif I to a serine (**Fig. S4B**), as approximately 62% of Def2 proteins. This further indicates that the A48S substitution does not impair PDF activity in *V. cholerae*.

**Fig. 4:**
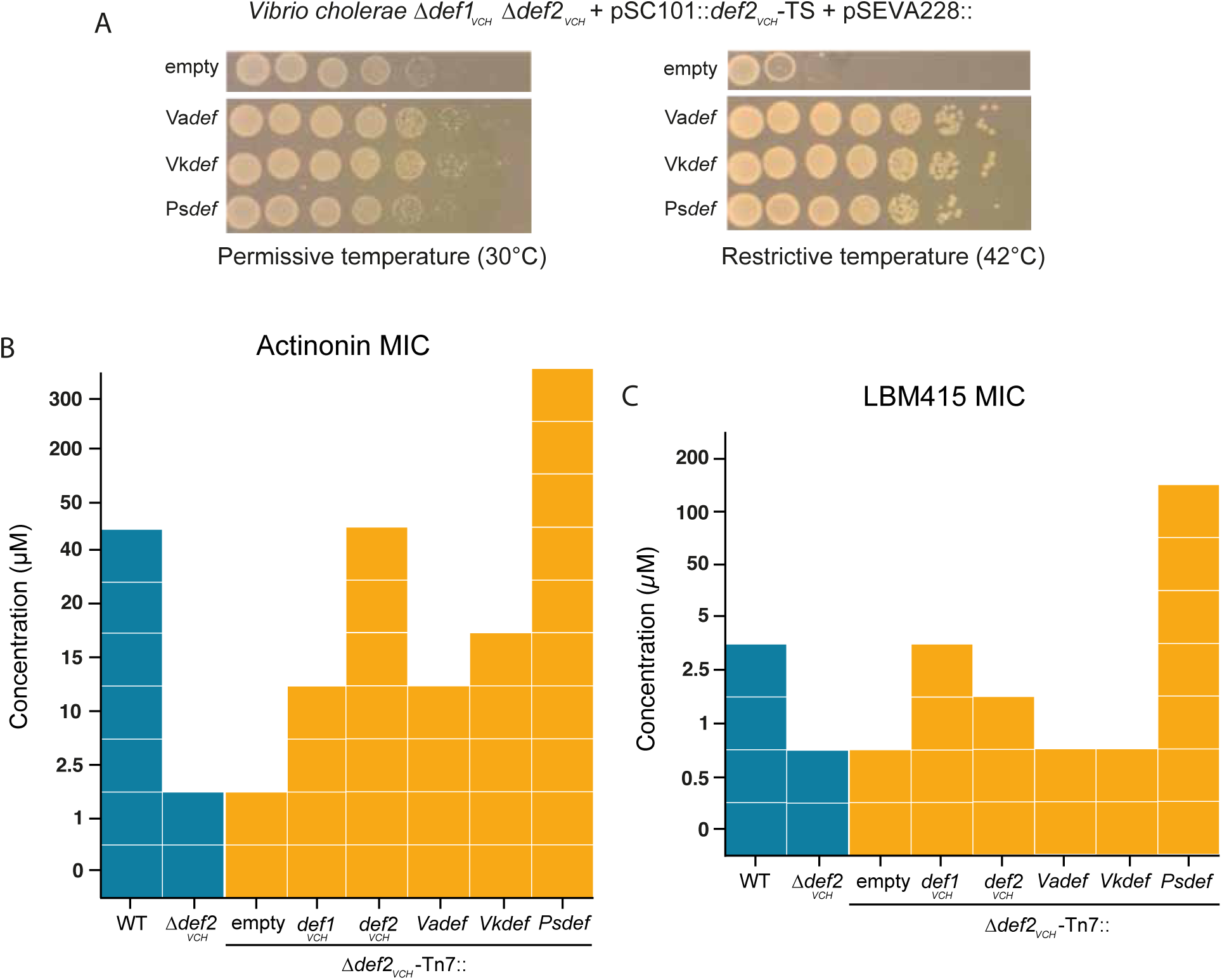
Activity and resistance profile of integron PDF. *A In vivo* PDF activity assay. Drop test of *V. cholerae* Δ*def1_VCH_* Δ*def2_VCH_* harboring a pSC101::*def2_VCH_*-TS plasmid and a pSEVA228 plasmid at permissive temperature (left panel) or restrictive temperature (right panel). Growth at restrictive temperature indicate that the PDF coding in the pSEVA228 plasmid is enzymatically active. VaDef, VkDef and PsDef correspond to *Vibrio anguillarum* str. MHK3 integron PDF, *Vibrio kanaloae* str. R17 integron PDF and *Pseudoxanthomonas suwonensis* str. 11-1 integron PDF respectively. **B)** Antibiotic susceptibility matrix of ACT. CMI was determine by broth microdilution as the minimal concentration of ACT with no observed growth. **C)** Antibiotic susceptibility matrix of LBM415. CMI was determine by broth microdilution as the minimal concentration of LBM415 with no observed growth.

According to structural predictions from AlphaFold3 <u>(Abramson *et al*., 2024)</u>, VaDef and VkDef, both belonging to the Def2 group, exhibit overall structure similarity with Def2_VCH_ (RMSD 0.38 Å over 151 alpha carbon and 0.39 Å over 153 alpha carbon, respectively). In contrast, PsDef, predicted to derive from a distinct evolutionary lineage (**Fig. 1E**), features an additional small C-terminal α-helix turn, typically absent in known type 1B PDFs (**Fig. S7A**).

### Integron PDF cassette confer resistance to PDF inhibitor

We assessed the resistance profiles of integron-encoded PDF cassettes VaDef, VkDef, and PsDef once integrated in single copy into the chromosome of the V*. cholerae* Δ*def2_VCH_* mutant, under the control of the constitutive *def2_VCH_* promoter. Unlike *V. cholerae* Def2_VCH_, the *Vibrio* integron cassettes VaDef and VkDef conferred only limited resistance to ACT and LBM415 in *V. cholerae* (**Fig. 4B-C**), despite also having a tyrosine at position 97. This limited resistance may be due to lower expression levels in *V. cholerae*, potentially resulting from differing codon usage. Consequently, these integron-encoded PDF cassettes might confer effective resistance to ACT in their native host.

In contrast, PsDef integration led to substantial resistance to actinonin, with at least a six-fold higher MIC compared to the WT strain and at least a 120-fold increase compared to the Δ*def2_VCH_* mutant. Growth was observed even at the highest concentrations tested, preventing a precise MIC determination (**Fig. 4B**). PsDef also conferred strong resistance to LBM415, resulting in a 40-fold MIC increase relative to the WT strain and a 200-fold increase compared to either the Δ*def1_VCH_* or Δ*def2_VCH_* mutants (**Fig. 4C**). Furthermore, expressing PsDef in *E. coli* also conferred resistance to peptide deformylase inhibitors, with the ACT MIC increasing three-fold and the LBM415 MIC increasing 25-fold. (**Fig. S7C-D**).

PsDef shares several characteristics with the Def2_VCH_ PDF of *V. cholerae*, notably the presence of tyrosine at position 97. It also features an insertion of a glycine between glutamates E87 and E88 of motif II (**Fig. S7B**). Glutamate E87 likely participates in ACT binding as observed in *E. coli* PDF and V*. cholerae* Def1_VCH_, while glutamate E88 is crucial for forming the hydrophobic pocket that accommodates the ACT P1’ pentane chain (Clements *et al*., 2001). This amino acid insertion may alter the spatial positioning of E87, hindering its interaction with ACT. Such structural modifications could decrease ACT binding affinity, potentially explaining the notable resistance PsDef confers to peptide deformylase inhibitors.

A significant risk associated with the PsDef cassette is its potential capture by a class 1 mobile integron, which could facilitate its transfer across different bacterial species. To evaluate this risk, we assessed the recombinogenic capacity of the cassette and found it to be considerable (**Fig. S8A-B**). Moreover, under the control of a native mobile integron Pc promoter, the expression of this cassette conferred significant resistance to both ACT and LBM415 in *V. cholerae* (**Fig. S8C-D**).

## Discussion

Despite the fact that the presence of accessory PDFs in certain bacterial genomes has been noted for over twenty years (Margolis *et al*., 2000; Guilloteau *et al*., 2002), no comprehensive study had yet addressed the global distribution and role of these accessory PDFs. We aimed at filling this gap and determined the distribution and genetic support of accessory PDFs and explored potential roles, which might explain their selection and maintenance in bacterial genomes.

Our large-scale analysis of complete genomes from RefSeq suggests that the formylation of initiator tRNA is likely conserved across most bacteria, as nearly all genomes encode a *fmt* gene. In addition, the presence of several PDF genes is a property common to ca. 50% of bacterial species. Additionally, we revealed significant intra-species variability for the presence of accessory PDFs, explained by their location on mobile genetic elements (MGEs) capable of horizontal gene transfer (HGT), such as plasmids and integrons. To our knowledge, PDF is the first essential gene found in integron cassettes. The presence of the exact chromosomal copy of *E. coli* PDF gene on plasmids in 12 *Enterobacteriaceae* species further demonstrated that PDFs could disseminate horizontally.

Until now, the role and activities of accessory PDFs had only been studied in a few species, notably *S. aureus* and *S. pneumoniae*, which possess an inactive type 3 accessory PDF (Margolis *et al*., 2000; Margolis *et al*., 2001), and *B. subtilis*, whose accessory PDF was shown to have similar enzymatic activity to the canonical PDF (Haas *et al*., 2001). Our analysis of reference genomes coding a single PDF allowed us to redefine the three conserved motifs previously described (Giglione *et al*., 2004). This analysis revealed that the vast majority of accessory PDFs retain the critical amino acids necessary for enzymatic activity. As 7% of bacterial species do not encode any PDF with all three motifs, this suggests that the precise presence of these motifs is not required for catalytic activity in all bacterial species. Additionally, earlier study has shown that type 3 PDFs of *T. brucei*, with mutations in these motifs, are active (Bouzaidi-Tiali *et al*., 2007), indicating that even the accessory PDFs lacking full conservation of these conserved motifs might have deformylase activity in certain ecological contexts, as seen with many eco-paralogs (Mazel and Marlière, 1989; Sanchez-Perez *et al*., 2008; Bratlie *et al*., 2010). Similarly to many proteins in hyperhalophilic bacteria which are active at high salinity levels, but exhibit minimal or no activity under standard salinity conditions (Sanchez-Perez *et al*., 2008), accessory type 3 PDFs, which exhibit low or no deformylase activity *in vitro*, might become active *in vivo* under specific environmental stresses. In this line, given that PDFs are metalloproteases and that two of the conserved motifs are involved in binding divalent metals, it is plausible that type 3 PDF enzymes might adapt to oxidative stress conditions. During oxidative stress, divalent cations can be oxidized to trivalent forms, potentially inactivating standard metalloproteases. Type 3 PDFs might bind alternative cofactors, such as non-divalent metals, enabling deformylase activity under oxidative stress. This activity, although potentially lower than that of canonical PDFs, could be sufficient to support bacterial survival and stress response.

Our comparative genomics analyses showed that many *Vibrionaceae* species encode multiple PDFs and most PDFs identified in integron cassettes are found in *Vibrio* genomes. Furthermore, the presence of two PDFs in *V. cholerae* was identified 20 years ago, but their roles and activities had not been studied yet. Our study revealed that about half of the *Vibrionaceae* species encode an accessory PDF, Def2, which belongs to the same phylogenetic group as PDFs found in *Vibrio* integron cassettes. Def2-like PDFs have strong structural homology with canonical Def1 PDFs but diverge at the C-terminal tail, involved in ribosome association. As, the ribosomal exit tunnel is particularly dynamic, with many co-translational proteins interacting near or overlapping the PDF binding site (Denks *et al*., 2017; Bhakta *et al*., 2019; Akbar *et al*., 2021; Bögeholz *et al*., 2021), an accessory PDF with different ribosome association dynamics could alter co-translational protein binding on the ribosome, affecting nascent peptide processing. Moreover, the formyl group on the initiator methionine was recognized as a N-terminal degradation signal (N-degron), regulating nascent protein quality control (Piatkov et al., 2015). Having multiple PDFs might enhance deformylation flexibility process, especially under stress conditions that affect ribosomal structure.

In *V. cholerae*, we showed that both PDFs, Def1_VCH_ and Def2_VCH_, are active and support bacterial growth in rich media. Accessory Def2_VCH_ confers greater resistance to ACT, a natural PDF inhibitor produced by Streptomyces bacteria, compared to the canonical Def1_VCH_. Characterization of both *V. cholerae* PDFs complexed with ACT revealed that Def2_VCH_ exhibits a lower affinity for ACT than Def1_VCH_, attributed to amino acid substitutions affecting ACT binding, thereby explaining the increased resistance observed. Given the phylogenetic similarity of integron PDF cassettes in Vibrio species to the Def2 group, we hypothesized that these cassettes might also confer resistance to ACT in *V. cholerae*. However, the *Vibrio* integron PDF cassettes tested conferred lower resistance than Def2_VCH_, despite sharing similar substitutions affecting ACT affinity, likely due to suboptimal codon usage.

It is unlikely that the presence of actinonin in the environment has driven the selection of accessory PDFs in *Vibrio*, given that *Streptomyces* and *Vibrio* do not occupy overlapping ecological niches. Instead, it is more plausible that other, yet unidentified, biosynthetic gene clusters (BGCs) producing PDF inhibitors exist within *Vibrio*’s natural habitat. One such BGC, previously identified in *Vibrio* species, has been shown to mediate bacterial competition in marine environments (Cordero *et al*., 2012; Costa *et al*., 2024), and recent studies confirm that it produces a peptide-based PDF inhibitor (Chen *et al.,* 2025; Rill *et al*., 2025). The presence of this BGC may therefore have contributed to the selection and maintenance of accessory PDFs in *Vibrio* genomes. Supporting this idea, another BGC producing a PDF inhibitor has also been identified in *Rhodococcus fascians*, with its co-encoded PDF conferring resistance to the compound (Ford et al., 2024). These findings suggest that naturally occurring PDF inhibitors are more widespread than previously thought, and that selective pressure imposed by such compounds may be a general driver for the acquisition of accessory PDFs in bacterial genomes.

Finally, the integron PDF cassette from *P. suwonensis* (PsDef) has been shown to confer high resistance to both ACT and LBM415, even when expressed under a native mobile integron promoter. Furthermore, this cassette’s high recombinogenic capability suggests it could be horizontally transmitted between bacterial species via plasmid-harboring mobile integrons. Due to its resistance phenotype, the dissemination of this cassette could pose a challenge for the therapeutic use of PDF inhibitors and question their viability in clinical applications.

Given that PDF inhibitors are likely present in bacterial environments, they probably contribute to bacterial competition. This environmental presence may have selected for PDFs resistant to these inhibitors, partly explaining the widespread occurrence of multiple PDF genes in bacterial genomes. However, the exact mechanisms driving the selection and maintenance of accessory PDF genes are likely multifaceted, involving resistance to natural inhibitors and other stresses, such as those influencing ribosome integrity.

## Acknowledgement

We would like to thank M.E. Val, J. Czarnecki, M. Lang and the Institut Pasteur strain collection (CIP) for providing several strains and/or plasmids. We would like to thank T. Niault for his thoughtful review of the manuscript. We thank H. Vaysset for his feedback on our phylogenetic analysis. We also thank the “Plate-forme de microbiologie mutualisée” (P2M), Pasteur International Bioresources Network (PIBnet), Institut Pasteur, Paris, France for their whole genome bacterial sequencing service. We thank the staff from the Crystallography platform at Institut Pasteur and the synchrotron source SOLEIL (Saint-Aubin, France) for granting access to the facility. We thank the staff of the beamlines Proxima 1 and Proxima 2A for their advice and assistance during X-ray data collections. We thank S. Rosario, S. Brûlé and P. England from the Molecular Biophysics Platform, Institut Pasteur, Paris, France, for the protein quality control analysis. We thank Novartis for given us the LBM415 compound.

