## Supplementary Figures for "Unraveling the prevalence and multifaceted roles of accessory peptide deformylases in bacterial adaptation and resistance"

### Supplementary Materials

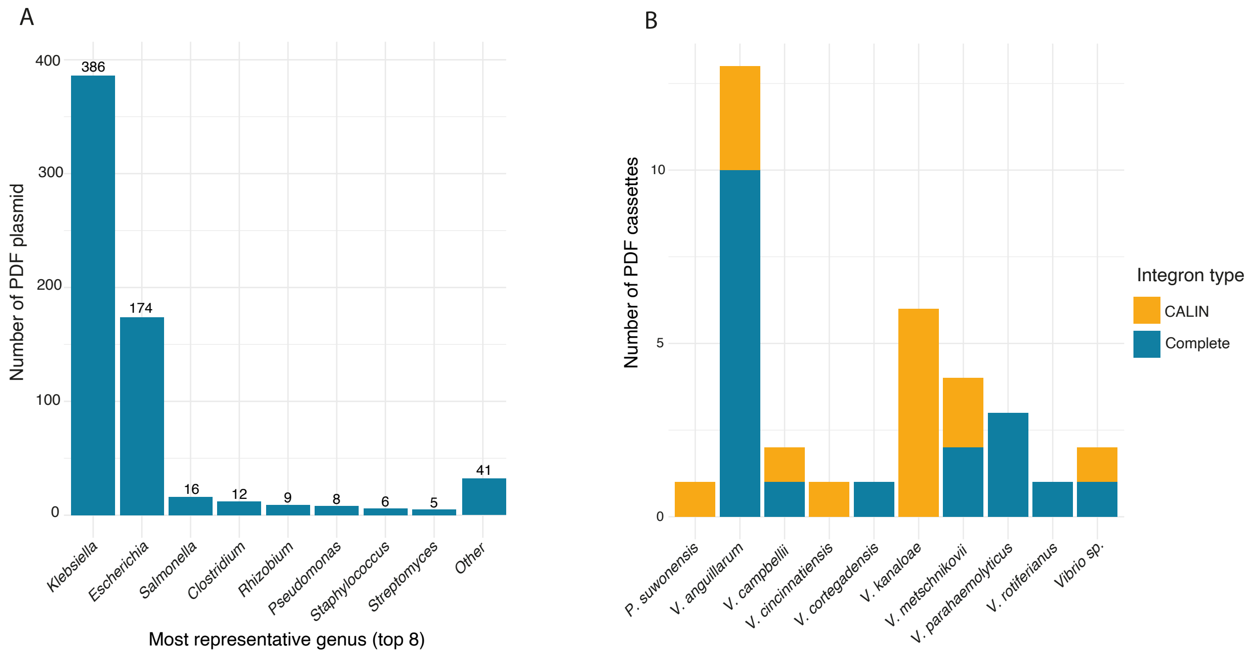


**Fig. S1: Distribution of PDF-encoding plasmids and integron cassettes. A)** Distribution of PDF-encoding plasmids across bacterial genera. Bars indicate the number of identified plasmids carrying PDF genes in each genus. Only the eight most represented genera are shown, with the corresponding counts displayed above the bars. **B)** Distribution of integron cassettes containing PDF genes across various bacterial species. Bars indicate the number of PDF cassettes, with colors distinguishing CALINs (yellow) from complete integrons (blue).

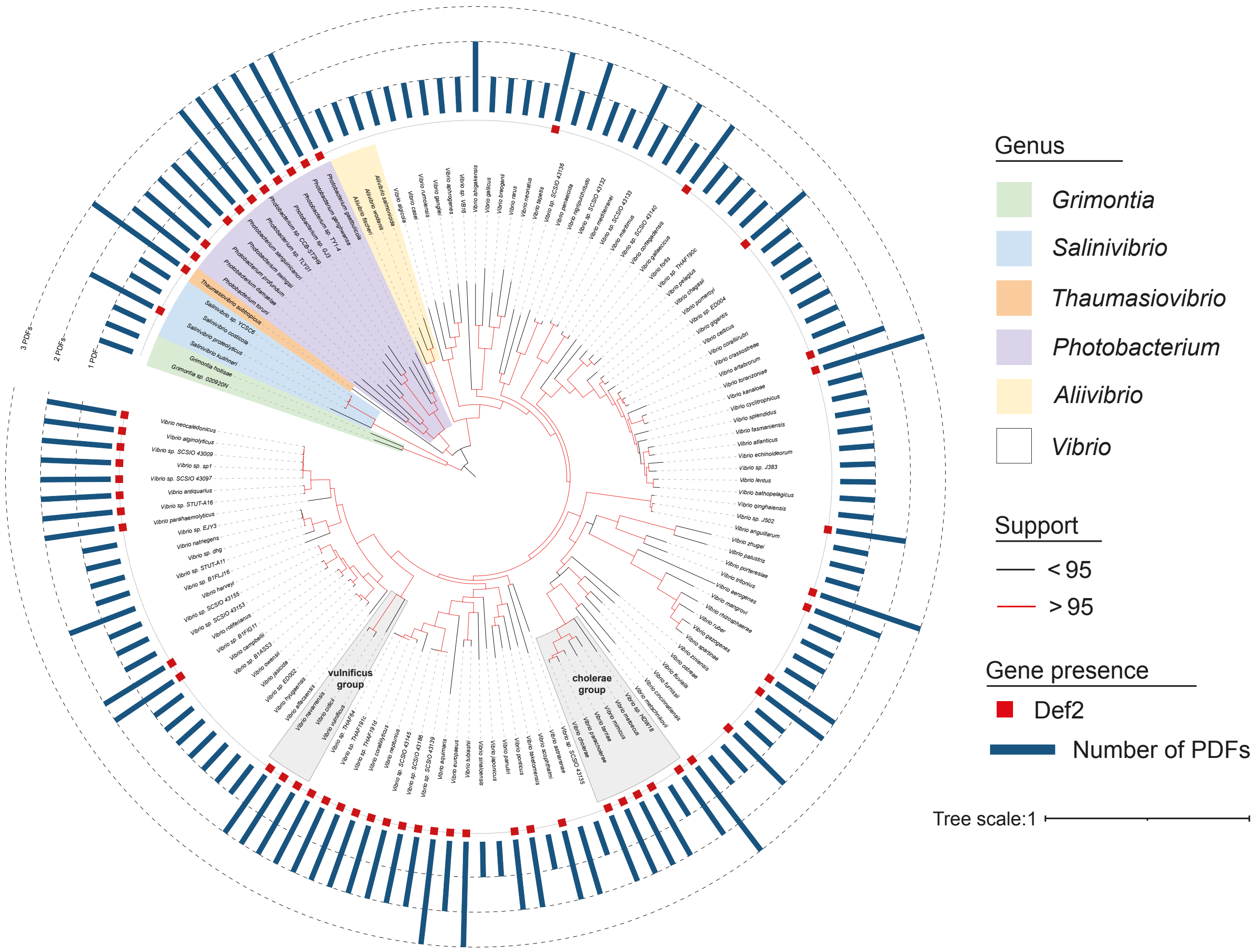


**Fig. S2: Phylogenetic tree of 136 complete *Vibrionaceae* species (Table S2).** The bar plot indicates the number of PDFs, with Def2 PDFs presence highlighted by red squares. The core genome was used for phylogenetic analysis. Genus groups are color-coded for clarity. Branches with an UFBoot support values greater than 95 are shown in red. The list of strain used for the phylogeny as well as the number of PDF per strain and the presence of Def2 protein are summarized in **Table S2**.

**Fig. S3: Phylogenetic tree of Vibrionaceae Def2 proteins.** The Def2 sequences used in these phylogenetic trees are from the species listed in Fig. S2 and Table S2. Sequences were aligned using either MUSCLE super5 (four first tree) or MAFFT (bottom tree) (Edgar, 2022; Katoh and Standley, 2013). Multiple Sequence Alignments (MSAs) generated by MUSCLE were created by permuting the guide tree ("none", "abc", "acb", "bca"). Branches with an UFBoot support values greater than 80 are shown in red. Trees were rooted using height Def1 sequences as an outgroup. This approach allows us to test the stability of the identified groups against MSA perturbations. Our analysis consistently identified the 'cholerae/vulnificus' group across all five trees, indicating a horizontal gene transfer event across these species.

**Fig. S3:**

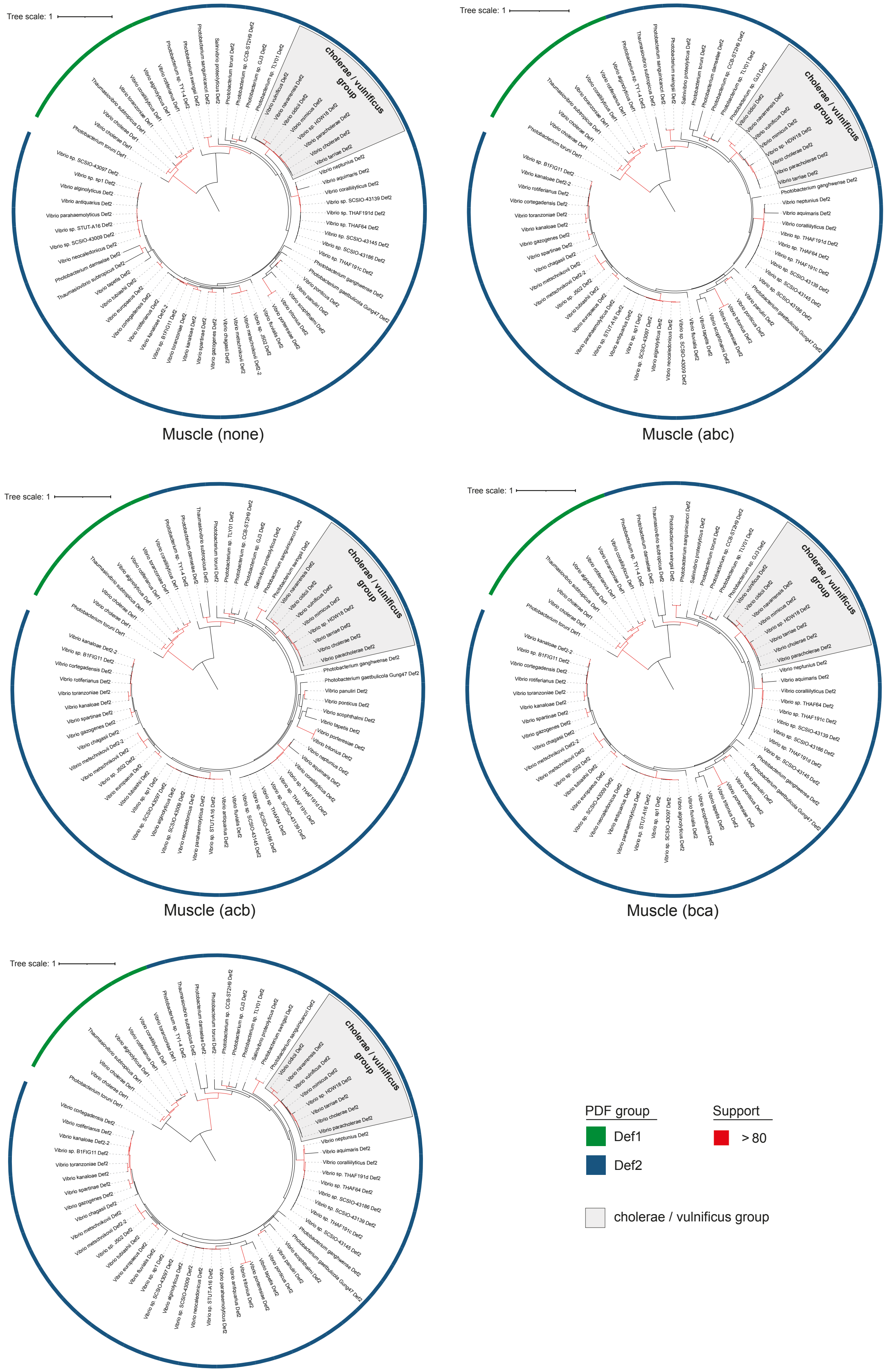


MAFFT L-INS-i

**Fig. S4: Structure and sequence comparison of *Vibrio cholerae* Def1$_{VCH}$ and Def2$_{VCH}$. A)** Structural alignment of *V. cholerae* Def1$_{VCH}$ (PDB ID: 3FWX - blue) and *V. cholerae* Def2$_{VCH}$ (PDB ID: 3QU1 - green), shown in front and side views, highlighting overall structural conservation and significant divergence in the C-terminal region**. B)** Alignment of amino acid sequences from *E. coli* Def, *V. cholerae* Def1$_{VCH}$ and Def2$_{VCH}$ and *V. tubiashii* BGC-encoded PDF (BGC-Def). Conserved motifs (I, II, III) are boxed. ACT-binding amino acids are highlighted in red. In BGC-Def, these are predicted based on homology with *E. coli* and *V. cholerae* PDFs. Amino acids forming the hydrophobic pocket for the ACT P1' pentane chain are marked by an asterisk (*). Ribosome-binding residues in *E. coli* are in green; these are putative in other PDFs, inferred from *E. coli* homology. Arginine at position 153 and lysine at 163 are absent in Def2$_{VCH}$ and BGC-Def, homologous residues are highlighted in lighter green. **C)** Conservation of the six canonical key amino acids involved in ribosome binding in *Vibrio* Def1 and Def2 group. **D)** Focus on the amino acids conservation at the position 153 and 163 in Def1 and Def2 group.

**Fig. S4:**

A

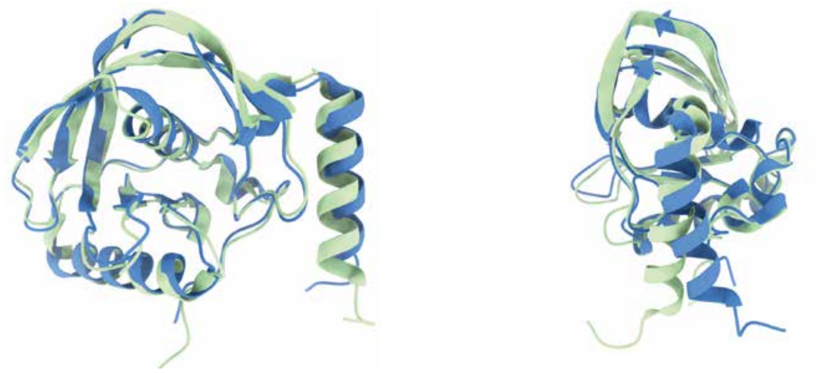

B

```
                        1          10         20         30         40    *    50
E. coli Def -           (M)SVLQVLHIP DERLRKVAKP VEEVNAEIQR IVDDMFETMY AEEGIGLAAT QVDIHQRIIV
V. cholerae Def1VCH-    (M)SVLQVLTFP DDRLRTVAKP VEQVTPEIQQ IVDDMLETMY AEEGIGLAAT QVDIHQRIVV
V. cholerae Def2VCH-    (M)AVLEILTAP DPRLRVQSKQ VTDVASVQTL IDDLLDTLYA TDNGIGLAAP QVGREEAIVV
V. tubiashii BGC-Def -  (M)AIRPIIEVP DDRLRVTYQI VDDVSTVQTL IDDMLDTLYD TDNGIGLAAA QIGHSEAVLI
                                                                          Motif I

                        60         70         80       * 90 *      100        110
E. coli Def -           IDVSENRDER LVLINPELLE KSGETGIEEG CLSIPEQRAL VPRAEKVKIR ALDRDGKPFE
V. cholerae Def1VCH-    IDISETRDQP MVLINPEIIE KRGEDGIEEG CLSVPGARAL VPRAAEVTVK ALDRNGQEYQ
V. cholerae Def2VCH-    IDLSDNRDQP LVLINPKVVS GSNKEMGQEG CLSVPDYYAD VERYTSVVVE ALDREGKPLR
V. tubiashii BGC-Def -  IDISENRDQP LIMINPQIIE HEGLIDSEEG CLSVPGVYAK VQRHQRVKVQ ALNRQGEAFT
                                                       Motif II

                        120      * 130 *    *  140        150        160
E. coli Def -           LEADGLLAIC IQHEMDHLVG KLFMDYLSPL KQQRIRQKVE KLDRLKARA
V. cholerae Def1VCH-    FDADDLLAIC VQHEMDHLAG KLFVDYLSPL KRNRIKEKLE KIKRFNEKK
V. cholerae Def2VCH-    IETSDFLAIV MQHEIDHLSG NLFIDYLSPL KQQMAMKKVK KHVKNRAR
V. tubiashii BGC-Def -  IEDDDYLAIV MQHEIDHLHG KIFIDYLSPL KRKMAMKKIK KFQRAKVSAA
                                   Motif III
```

C

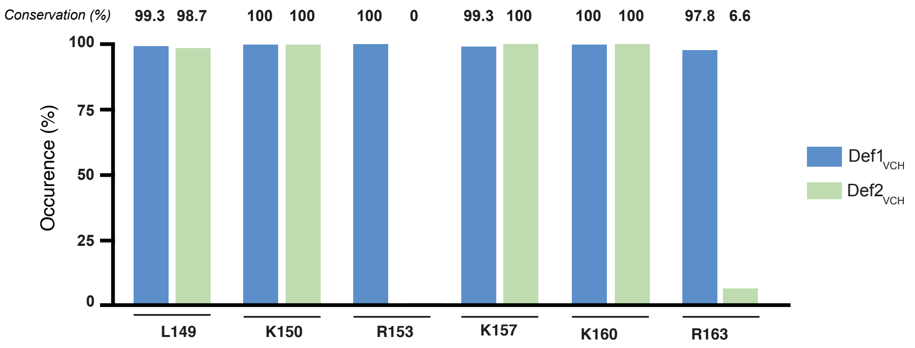


D

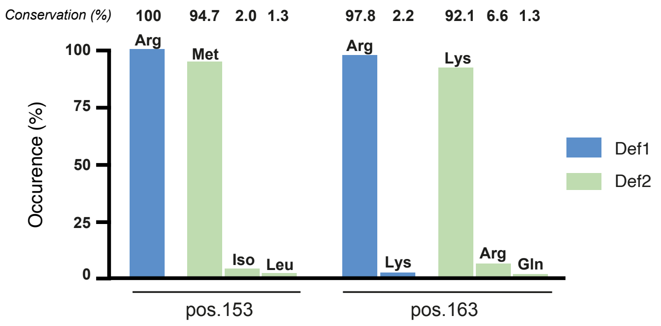

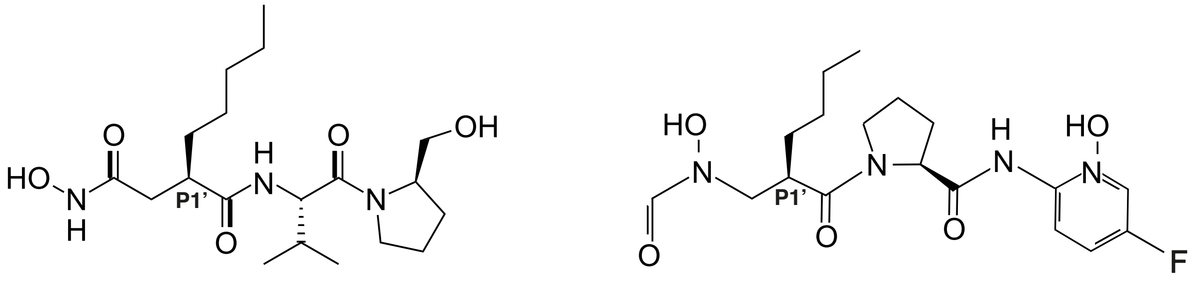

Actinonin LBM415

**Fig. S5: Chemical structure of actinonin and LBM415.**

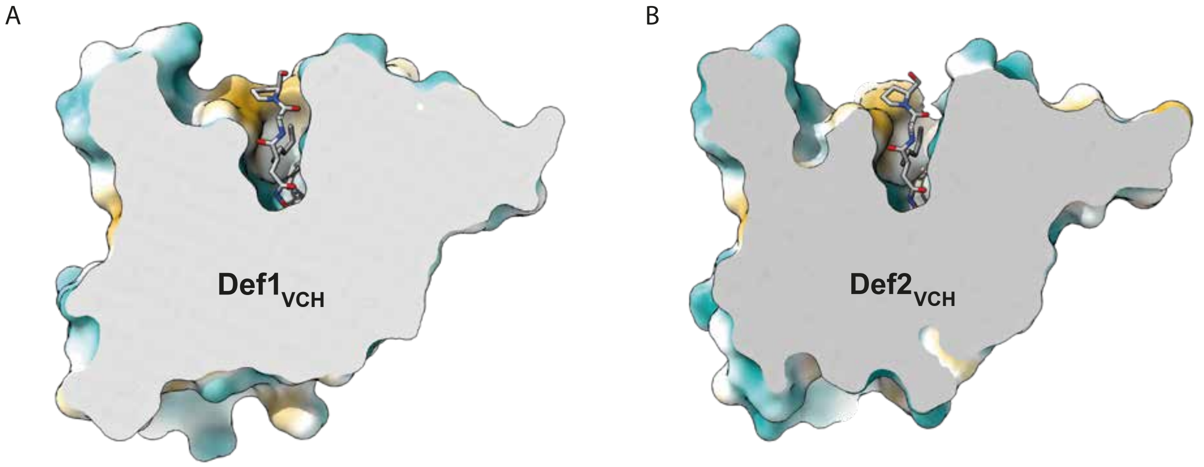


**Fig. S6: Structural view of actinonin embedded in the catalytic cavities of V. cholerae PDFs.** (A) Crystal structure of Def1$_{VCH}$ and (B) Def2$_{VCH}$ in complex with ACT, shown in stick representation. The surface view highlights the insertion of ACT into the catalytic pocket of both enzymes.

**Fig. S7: Structure and *E. coli* Δ*tolC* resistance profile of the integron-encoded PDF of *P. suwonensis* 11-1. A)** Structural alignment of *E. coli* Def (PDB ID: 1DFF – green) and PsDef (AlphaFold3 prediction – blue), shown in front and side views, highlighting overall structural conservation of classical type 1B PDF and the presenece of an additional C-terminal α helix**. B)** Alignment of amino acid sequences from *E. coli* Def and *P. suwonensis* integron-encoded PDF (PsDef). Conserved motifs (I, II, III) are boxed. ACT-binding amino acids are highlighted in red. In PsDef, these are predicted based on homology with *E. coli* and *V. cholerae* PDFs. PsDef's glutamate E90 is highlighted in orange due to a glycine insertion that may affect its interaction with ACT. Amino acids forming the hydrophobic pocket for the ACT P1' pentane chain are marked by an asterisk (*). Ribosome-binding residues in *E. coli* are in green and are putative in PsDef, inferred from *E. coli* homology. Arginine at position 157 is absent in PsDef, homologous residue is highlighted in lighter green**. D**ue to the presence of four amino acid insertions in PsDef, the alignment numbering differs from the canonical *E. coli* Def sequence. In the main amino acid positions refer to the *E. coli* Def sequence. **C-D)** Antibiotic susceptibility matrix of *E. coli ΔtolC* strain harboring low copy number plasmid pSC101 expression either *E. coli* PDF (EcDef), Def1$_{VCH}$, Def2$_{VCH}$ or PsDef, for ACT (C) and LBM415 (D).

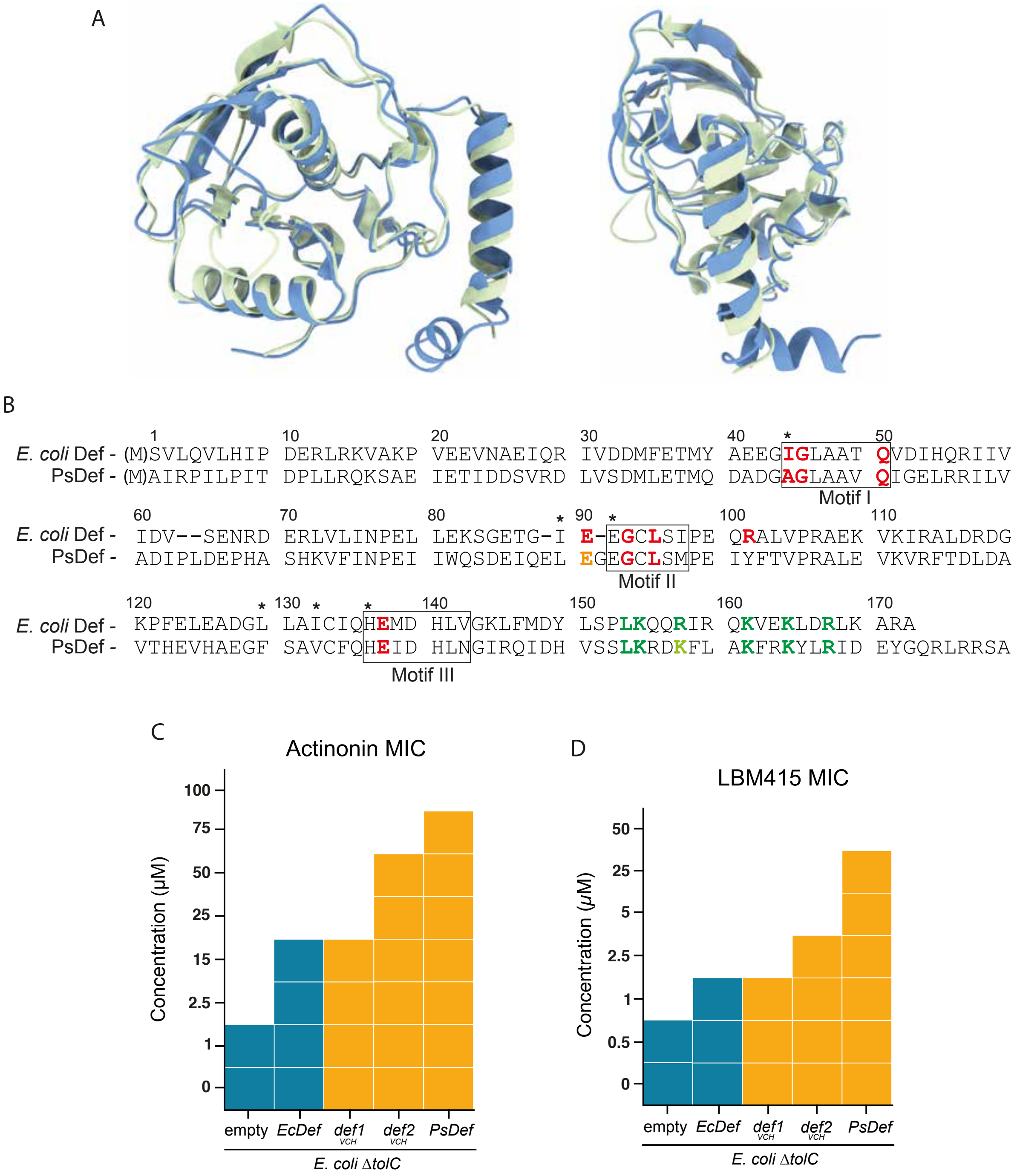
A
B
1 10 20 30 40 * 50
E. coli Def - (M)SVLQVLHIP DERLRKVAKP VEEVNAEIQR IVDDMFETMY AEEGIGLAAT QVDIHQRIIV
PsDef - (M)AIRPILPIT DPLLRQKSAE IETIDDSVRD LVSDMLETMQ DADGAGLAAV QIGELRRILV
Motif I
60 70 80 * 90 * 100 110
E. coli Def - IDV--SENRD ERLVLINPEL LEKSGETG-I E-EGCLSIPE QRALVPRAEK VKIRALDRDG
PsDef - ADIPLDEPHA SHKVFINPEI IWQSDEIQEL EGEGCLSMPE IYFTVPRALE VKVRFTDLDA
Motif II
120 * 130 * * 140 150 160 170
E. coli Def - KPFELEADGL LAICIQHEMD HLVGKLFMDY LSPLKQQRIR QKVEKLDRLK ARA
PsDef - VTHEVHAEGF SAVCFQHEID HLNGIRQIDH VSSLKRDKFL AKFRKYLRID EYGQRLRRSA
Motif III
C
Actinonin MIC
Concentration (µM)
100
75
50
25
15
2.5
1
0
empty
EcDef
def1 VCH
def2 VCH
PsDef
E. coli ΔtolC
D
LBM415 MIC
Concentration (µM)
50
25
5
2.5
1
0.5
0
empty
EcDef
def1 VCH
def2 VCH
PsDef
E. coli ΔtolC

**Fig. S8: Resistance and recombinogenic potential of the *psDef* cassette in a natural context.** **A)** Structure of the recombinogenic $attC_{psDef}$. **B)** Recombination rate of the *attI*1 x $attC_{psDef}$ and *attI*1 x $attC_{aadA7}$ reactions. * Indicate that the recombination rate was below the detection limit. **C-D)** Antibiotic susceptibility matrix of *psDef* on a pSC101 plasmid under variable natural Pc promoter for ACT (C) and LBM415 (D). Grey boxes indicated spontaneous apparition of a P2 promoter during the assay, conferring an increased resistance profil (see extended figure S7).

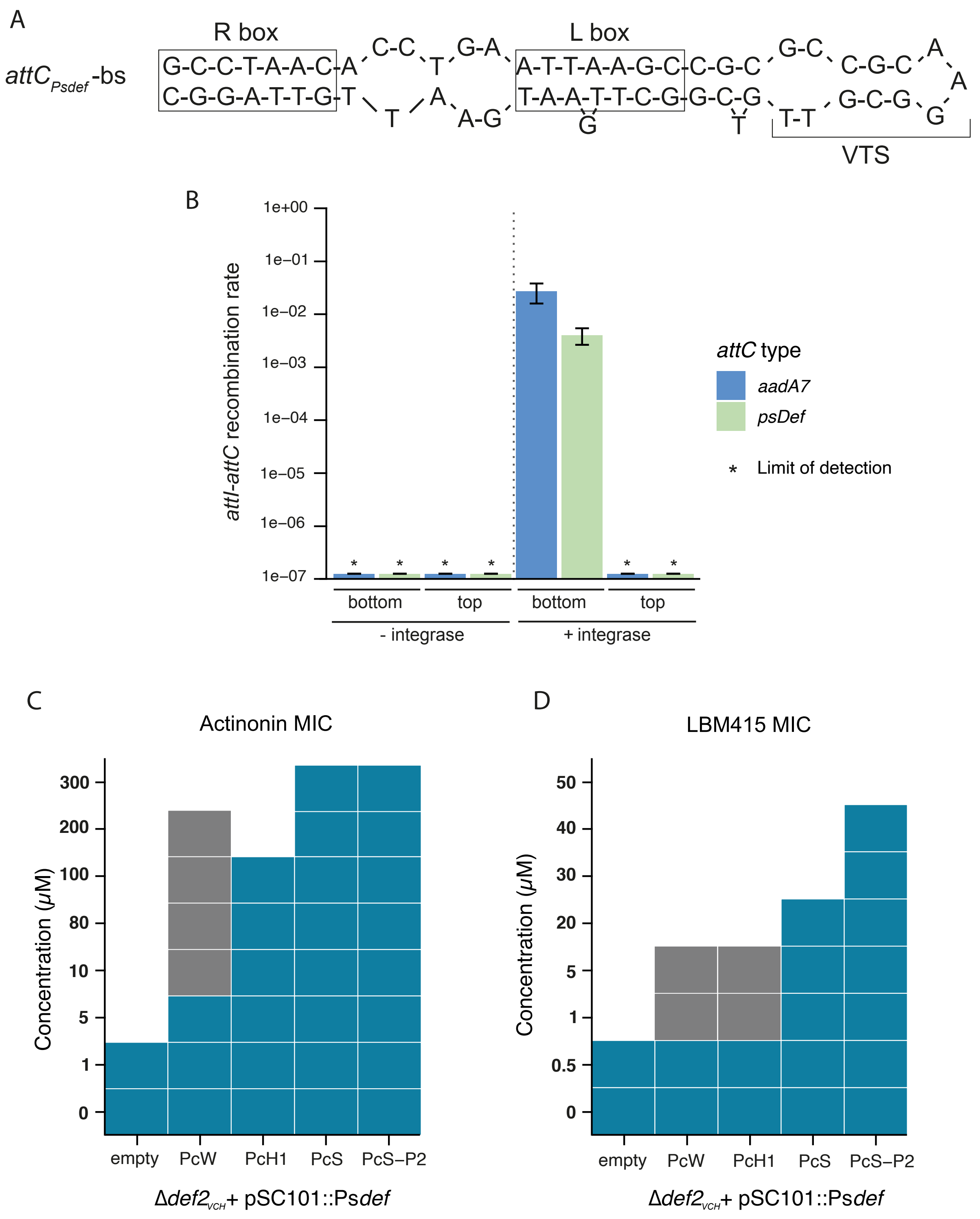
A
attC Psdef -bs
R box
L box
VTS
B
attI-attC recombination rate
1e+00
1e−01
1e−02
1e−03
1e−04
1e−05
1e−06
1e−07
bottom
top
- integrase
+ integrase
attC type
aadA7
psDef
* Limit of detection
C
Actinonin MIC
Concentration (µM)
300
200
100
80
10
5
1
0
empty
PcW
PcH1
PcS
PcS−P2
Δdef2 VCH + pSC101::Psdef
D
LBM415 MIC
50
40
30
20
5
1
0.5
0

**Extended data – Fig. S8:**

A significant risk associated with the PsDef cassette lies in its potential capture by a class I mobile integron, facilitating its transfer across various bacterial species and potentially driving resistance to PDF inhibitors.

To assess whether the PsDef cassette confers resistance to PDF inhibitors when integrated and expressed within a mobile integron, we cloned the PsDef cassette into a low copy number plasmid under the control of three different native integron promoters: PcW, PcH1, and PcS, which represent 94% of the Pc promoters found in mobile integrons, as well as a P2 promoter, present in 8.7% of integrons (Jové *et al.*, 2010). This experimental setup aimed to simulate the natural integration of the cassette into a class 1 mobile integron, an event that could occur in natural environments. The resistance to PDF inhibitors conferred by the PsDef cassette in *V. cholerae* depended on the expression level driven by these promoters, as expected. The highest resistance levels were observed under the PcS promoter, which matched the resistance levels seen when the cassette was integrated into the chromosome. Under the PcH1 promoter, the resistance conferred was lower but still represented a 40-fold increase compared to the Δ$def2_{VCH}$ mutant and a two-fold increase compared to the WT strain. Conversely, under the PcW promoter, the resistance levels were significantly lower, with a two-fold increase over the Δ$def2_{VCH}$ mutant. Notably, we observed variability in resistance levels conferred by the PsDef cassette under the PcW promoter across replicates, with MIC values ranging from 5 µM to 200 µM. This suggested the rapid emergence of compensatory mutants. Upon analysis, we identified the spontaneous and rapid emergence of a P2 promoter (Collis and Hall, 1995; Jové *et al.*, 2010), which significantly increased PsDef expression and led to a 20-fold increase in resistance compared to the cassette under the PcW promoter alone (**Fig. S8C**). Similar results were obtained with LBM415, although the resistance levels were generally lower than those observed with actinonin (**Fig. S8D**).

We also investigated the recombinogenic capacity of the *psDef* cassette. To do this, we calculated the pFold value of its *attC* site, which reflects the inherent probability of the site forming a recombinogenic structure (**Fig. S8A**) (Loot *et al.*, 2017). The pFold value for the *psDef* cassette was determined to be 1, indicating a theoretically high recombinogenic potential. To confirm this, we measured the frequency of the *attI1* x *attC* reaction, which reflect the integration efficiency of the cassette into the first position of an integron. The *attI1* x $attC_{PsDef}$ recombination frequencies were found to be similar to those of the *attI1* x $attC_{aadA7}$, which is representative of highly recombinogenic *attC* sites (**Fig. S8B**). These results validate the high recombinogenic nature of the *psDef* cassette and its significant capacity to be mobilized by integron integrases.

#### Supplementary Table:

**Table S1: Strains and plasmids used in the study.**

**Table S2: List of the 136 Vibrionaceae genomes used for *Vibrionaceae* phylogeny.** For each genome, the RefSeq Genome Assembly identifier, the name of the Species as well as the number of PDF genes encoded and the presence of a PDF belonging to the Def2 clade are shown in the table.

**Table S3: List of genome not encoding for any PDF gene.** The genome size as well as the number of CDSs and pseudogenes encoded in the genome is shown in the table. In general, endosymbiotic bacteria have a small genome encoding only a limited number of genes. However, in this list, 2 genomes, *Otariodibacter oris* and *Tardibacter chloracetimidivorans*, are not endosymbiotic and also lacked PDF genes, as well as the methionyl-tRNA-formyltransferase *fmt*. For *O. oris*, another genome, though incomplete, contains both *def* and *fmt* genes, suggesting that the reference genome might be incomplete as well. For *T. chloracetimidivorans*, since no other genome of this species is available, it is difficult to determine whether this bacterium initiates translation without a formyl-methionine or if its deposited genome is incomplete.

**Table S4: Plasmids encoding a peptide deformylase (PDF) gene.** For each plasmid, the accession number, replicon name, size, and host organism are listed. The table also indicates the number of *def* genes encoded per replicon and its predicted mobility type (pCONJ: conjugative; pMOB: mobilizable; pdCONJ: decayed conjugative; pOriT: plasmid carrying only an origin of transfer; pMOBless: lacking known mobility markers). Supplementary sheets summarize the distribution of PDF-encoding plasmids across genera and species. For each taxon, the total number of plasmids in the database, the number encoding a *def* gene, and the corresponding percentage are provided, highlighting taxa where plasmid-borne PDFs are most frequent.

**Table S5: Genomes encoding a PDF gene within an integron.** For each genome, the RefSeq assembly ID, organism name, number of PDF-containing cassettes, integron type (complete integron or CALIN—*Cluster of attC sites lacking an integron-integrase*), total number of cassettes within the integron, position of the PDF cassette, and predicted integron mobility type are indicated.

**Table S6: Data collection and refinement statistics of the PDF-ACT crystallization experiment.**
